# *N*-acetyl-cysteinylated streptophenazines from *Streptomyces*

**DOI:** 10.1101/2021.12.06.470720

**Authors:** Kristiina Vind, Sonia Maffioli, Blanca Fernandez Ciruelos, Valentin Waschulin, Cristina Brunati, Matteo Simone, Margherita Sosio, Stefano Donadio

**Affiliations:** NAICONS Srl, 20139 Milan, Italy; Host-Microbe Interactomics Group, Wageningen University, 6708 WD Wageningen, The Netherlands; School of Life Sciences, University of Warwick, Coventry, United Kingdom

## Abstract

Here, we describe two *N*-acetyl-cysteinylated streptophenazines (**1** and **2**) produced by soil-derived *Streptomyces* sp. ID63040 and identified through a metabolomic approach. These metabolites attracted our interest due to their low occurrence frequency in a large library of fermentation broth extracts and their consistent presence in biological replicates of the producer strain. The compounds were found to possess broad-spectrum antibacterial activity while exhibiting low cytotoxicity. The biosynthetic gene cluster from *Streptomyces* sp. ID63040 was found to be highly similar to the streptophenazine reference cluster in the MIBiG database, which originates from the marine *Streptomyces* sp. CNB-091. Compounds **1** and **2** were the main streptophenazine products from *Streptomyces* sp. ID63040 at all cultivation times, but were not detected in *Streptomyces* sp. CNB-091. The lack of obvious candidates for cysteinylation in the *Streptomyces* sp. ID63040 biosynthetic gene cluster suggests that the *N*-acetyl-cysteine moiety derives from cellular functions, most likely from mycothiol. Overall, our data represent an interesting example on how to leverage metabolomics for the discovery of new natural products and point out to the often-neglected contribution of house-keeping cellular functions to natural product diversification.

**Graphical abstract:** 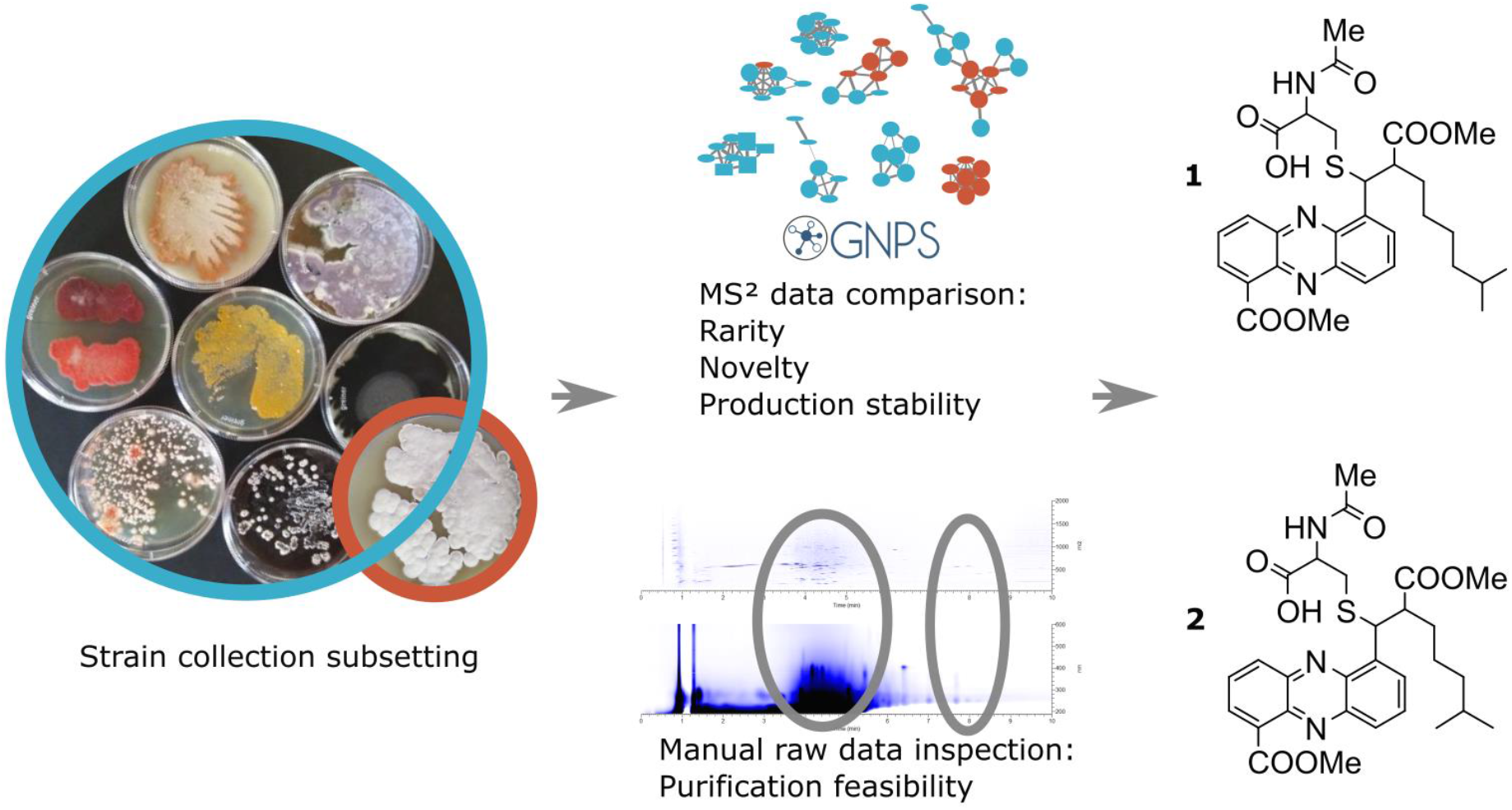

## INTRODUCTION

Resistance has been observed against all established classes of clinically relevant antibiotics^1^, rendering once easy-to-cure diseases difficult to treat. Hence, there is an urgent need for novel chemical scaffolds with antibacterial activity. With synthetic approaches performing below expectations, natural products are still the prevalent source of antibiotics in human use^2^. Bioactivity based screenings introduce bias into natural products discovery, highlighting mostly the metabolites which are both frequently encountered and present in concentrations above the bioactivity test threshold^3–5^. To expand our knowledge of the vast chemical space covered by natural products, it is necessary to find alternative strategies to identify novel compounds to meet our medical needs.

One of the main challenges in natural products discovery lies in avoiding re-discovery of known scaffolds. The widespread use of mass spectrometry and rapid development of spectrum processing and analysis tools^6–9^ has facilitated recognizing previously described compounds in a process known as dereplication. As databases are biased towards bioactive metabolites and only a fraction of the occurring metabolites have been annotated, only a minority of signals in the metabolic fingerprint of any microbe can be dereplicated automatically. This implies that potential novelty of an unknown metabolite cannot be deduced from absence of annotation.

Relying on the assumption that novel chemistry is detected relatively rarely when exploring well-established microbial taxa^3^, it should nevertheless be possible to pinpoint medically valuable metabolites by exploring a sufficiently large dataset of metabolites. In this respect, NAICONS’ metabolic fingerprint library (NMFL), derived from about 14 000 extracts obtained from about four thousand actinomycete strains^10^, offers a great opportunity for “rarity-based” prioritization of metabolites to discover novel antimicrobials. Recently, a metabolomics-guided approach has enabled the discovery of the unusual biaryl-linked tripeptides produced by some *Planomonospora* strains and permitted the finding that the associated biosynthetic gene clusters (BGCs) are widespread among actinobacteria, although the corresponding metabolites have so far escaped detection^11^.

We describe the discovery of reliably produced antimicrobial streptophenazines featuring an *N*-acetylated cysteine attached to the alkyl chain C1’ via a thioether bridge through untargeted metabolomics^12^ approach. We trace the abundance dynamics of these molecules and compare the BGC from the producer strain *Streptomyces* sp. ID63040 to the streptophenazine reference cluster^13^.

## RESULTS AND DISCUSSION

### Metabolite prioritization

Current results stem from a study in aimed at identifying small molecule elicitors that would trigger production of secondary metabolites. Using 21 randomly chosen *Streptomyces* strains from the NAICONS’ collection of 45 000 actinomycete strains^10^, we cultivated them in medium-scale liquid cultures in a single medium but in separate experiments and analyzed the metabolite fingerprints of the corresponding extracts by LC-MS/MS (K.V., unpublished results). During the course of this study, we realized that the available dataset could be “mined” for metabolites that fulfilled the following properties: i) they were not detected in any of the 14 000 metabolic fingerprints contained in the NMFL; ii) they did not cluster together with any of the annotated metabolites in the GNPS platform^6^; iii) they co-eluted with few other metabolites in reverse phase HPLC; and iv) they were observed in extracts obtained from biological replicates of the same strain performed at different times (Figure 1).

**Figure 1.**
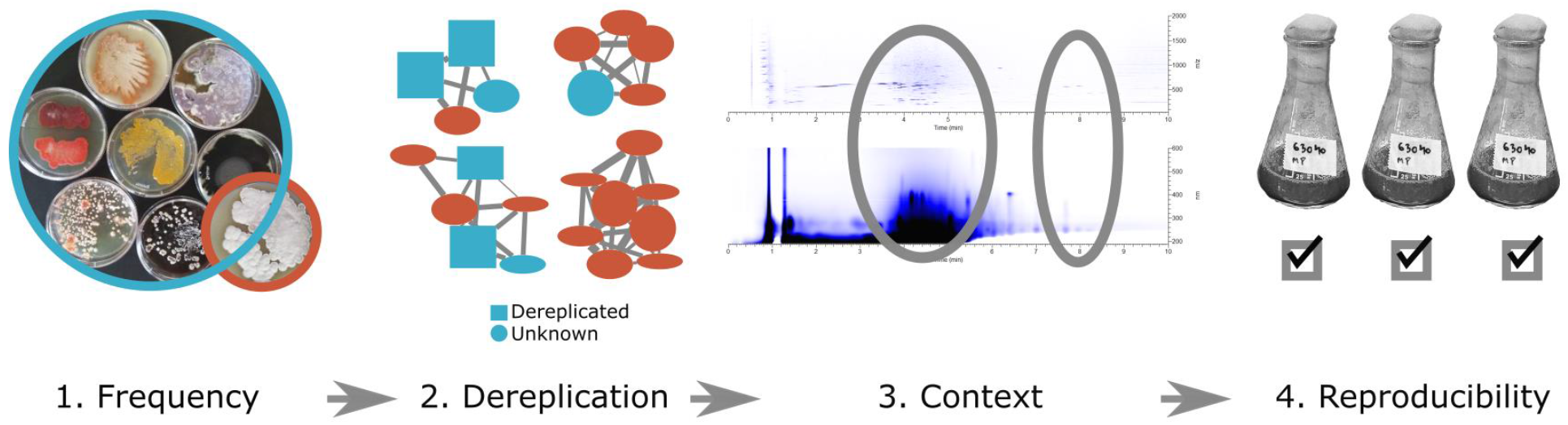
Metabolite prioritization workflow. First, we selected signals absent in the NMFL dataset (Frequency); next, we excluded signals associated with annotated molecular families in the GNPS database (Dereplication); then, we manually examined raw data to establish chromatogram crowdedness (Context); and finally, we verified the presence of signals across biological replicates (Reproducibility). We used GNPS for steps 1, 2 and 4, but in principle, any other suitable software of choice can be used, and the order of steps 3 and 4 can be reversed.

The metabolites occurring in the dataset related to 21 strains and absent in NMFL were considered as rare and potentially novel. The molecular network of 11362 datafiles related to roughly 4000 strains and 42 datafiles related to the set of 21 strains was calculated on GNPS server (K.V., unpublished results) The resulting table was analyzed with R as follows. First, we omitted all single nodes, as the presence of structurally related congeners strengthens metabolite identification. Then, we focused on nodes encountered only in the dataset of 21 strains. Next, we eliminated signals clustering into molecular families with characterized compounds. This logical flow, which was designed for identification of rarely occurring metabolites that were structurally unrelated to known molecules, led to the identification of 18 molecular families worthy of further investigation.

In the absence of other prioritization criteria, context assessment is a key step for increasing the odds for successful structural characterization. The metabolic fingerprint of a given strain is usually a complex matrix in which not all metabolites can be easily purified due to co-occurrence with structurally unrelated compounds with similar charge, polarity, and/or size. Obtaining enough compound with reasonable purity can be a rate-limiting step in natural product discovery that may require scaling up fermentation and establishing tailor-made purification protocols^14^. To validate our approach, we gave higher priority to compounds which appeared easy to purify on the basis of the crowdedness of the relevant portion of the reverse phase chromatogram. Furthermore, in the absence of co-eluting compounds, it is possible to unambiguously assign a UV-vis absorption spectrum to the metabolite of interest, which facilitates compound detection during purification. Manual inspection of raw data resulted in prioritization of just two molecular families from two strains.

Finally, to be able to carry out compound characterization using the natural host, it is critical to evaluate the production stability before deciding on which compound to pursue. The metabolites which were produced across experiments (biological replicates with different preparation of the same cultivation medium) were believed to be the result of a robust production process. Both molecular families prioritized in the previous step were present across experiments, representing good starting points for natural products discovery.

### Isolation and structure elucidation of N-acetyl-cysteinylated streptophenazines

The metabolites characterized in this paper were obtained from *Streptomyces* strain ID63040. Using LC-MS, we detected two main congeners in the lipophilic part of the chromatogram: *m/z* 584.2430 [M + H]^+^ (**1**) and 570.2277 [M + H]^+^ (**2**). Upon fragmentation, we observed neutral losses of 131.0036 Da (for **1**) and 131.0042 Da (for **2**), corresponding to C_4_H_5_N_1_O_2_S (calculated 131.0041 Da), thus establishing the presence of sulfur (Supplementary Table 1). The calculated molecular formulae for **1** and **2,** corresponding to the observed 584.2430 (calculated 584.2430) and observed 570.2277 (calculated 570.2274) [M + H]^+^, were hence C_30_H_35_N_3_O_7_S and C_29_H_37_N_3_O_7_S, respectively.

According to the available records, strain ID63040 was isolated from a soil sample collected near Ziniare, Burkina Faso on 06.06.1992. Based on its 16S rRNA gene sequence the closest relative is *Streptomyces cellostaticus* NBRC 12849, with sequence identity of 99.44%.

Strain ID63040 was grown in 2.5L liquid production medium for three days. The culture broth was filtered through Whatman paper followed by EtOH extraction of the mycelial cake and MPLC fractionation, yielding 3.8 mg of **1** as a yellowish-brown, amorphous solid, with UV-vis maxima at 252 and 366 nm (Supplementary Figure 1).

NMR analysis of **1** in acetone-d_6_ established the presence of an *N*-acetylated cysteine residue (Table 1). Moreover, the characteristic UV spectrum and the presence of numerous aromatic carbons with chemical shifts around δ 141 and 143 ppm, suggested that **1** contains a phenazine core. Three hydrogens were detected in each of the aromatic rings. One of the remaining carbons was clearly connected to a carboxylic ester. Additionally, we detected three discrete spin systems: a branched aliphatic chain, one free carboxylic acid and an additional carboxylic ester (Table 1).

**Table 1.** NMR correlations of compounds **1** and **2**.

|  | **1** |  |  |  |  | **2** |  |
| --- | --- | --- | --- | --- | --- | --- | --- |
|  | acetone-d<sub>6</sub> at 330K |  | dmsO-d<sub>6</sub> at 350K |  |  | acetone-d<sub>6</sub> at 330K |  |
|  | 1H | 13C | 1H | 13C |  | 1H | 13C |
| 1' | 5.64 | missing | 5.42 | 45.9 | 1' | 5.64 | missing |
| 2' | 3.50 | 51.0 | 3.46 | 51.1 | 2' | 3.50 | 51.0 |
| 3'_a | 1.50 | 31.4 | 1.45 | 31.3 | 3'_a | 1.50 | 31.4 |
| 3'_b | 1.20 | 31.4 | 1.20 | 31.3 | 3'_b | 1.20 | 31.4 |
| 4' | 1.34 | 29.2 | 1.12 | 24.3 |  |  |  |
| 5' | 1.18 | 26.9 | 1.11 | 26.9 | 4' | 1.18 | 26.9 |
| 6' | 0.90 | 38.2 | 0.86 | 38.3 | 5' | 0.90 | 38.2 |
| 7' | 1.26 | 27.3 | 1.26 | 27.5 | 6' | 1.26 | 27.3 |
| 8' e 9' Me<sub>2</sub> | 0.67 | 21.8 | 0.65 | 22.7 | 7' and 8' | 0.67 | 21.8 |
| Aliph_COOMe | 3.81 | 51.2 | 3.70 | 51.7 | Aliph_COOMe | 3.81 | 51.2 |
| Aliph_COOMe | - | 175.0 | - | 174.5 | Aliph_COOMe | - | 175.0 |
| 1''_CysCOOH | - | 171.1 | - | 172.1 | 1''_CysCOOH | - | 171.1 |
| 2''_Cys_alpha | 4.80 | 52.2 | 4.37 | 52.3 | 2''_Cys_alpha | 4.80 | 52.2 |
| 3''_Cys_beta1 | 2.96 | 33.5 | 2.82 | 34.2 | 3''_Cys_beta1 | 2.96 | 33.5 |
| 3''_Cys_beta2 | 2.84 | 33.5 | 2.70 | 34.2 | 3''_Cys_beta2 | 2.84 | 33.5 |
| 4'' (NH) | 7.40 | - | 7.85 | - | 4'' (NH) | 7.40 | - |
| 5''_AcCys | - | 169.2 | - | 169.3 | 5''_AcCys | - | 169.2 |
| 6''_Cys_Me | 2.05 | 22.0 | 1.80 | 22.7 | 6''_Cys_Me | 2.05 | 22.0 |
| Ar_1 | - | 132.0 | - | 132.0 | Ar_1 | - | 132.0 |
| Ar_6 | - | 132.0 | - | 132.0 | Ar_6 | - | 132.0 |
| Ar_2 | 8.24 | 131.3 | 8.22 | 131.8 | Ar_2 | 8.24 | 131.3 |
| Ar_3 | 8.03 | 129.7 | 8.03 | 130.3 | Ar_3 | 8.03 | 129.7 |
| Ar_4 | 8.57 | 133.1 | 8.42 | 133.2 | Ar_4 | 8.57 | 133.1 |
| Ar_7 | 8.07 | 129.8 | 8.07 | 130.3 | Ar_7 | 8.07 | 129.8 |
| Ar_8 | 8.03 | 131.2 | 7.99 | 131.7 | Ar_8 | 8.03 | 131.2 |
| Ar_9 | 8.19 | 129.0 | 8.14 | 129.2 | Ar_9 | 8.19 | 129.0 |
| 4a | - | 142.0 | - | 141.5 | 4a | - | 142.0 |
| 5a | - | 141.0 | - | 140.0 | 5a | - | 141.0 |
| 9a | - | 143.0 | - | 143.3 | 9a | - | 143.0 |
| 10a | - | 141.0 | - | 141.5 | 10a | - | 141.0 |
| Arom_COOMe | 4.04 | 51.8 | 4.02 | 52.7 | Arom_COOMe | 4.04 | 51.8 |
| Ar-COOMe | - | 166.8 | - | 167.1 | Ar-COOMe | - | 166.8 |

The number of methyl esters was confirmed by basic hydrolysis with NaOH revealing one readily hydrolyzed methyl ester (5 min at room temperature) and a second that required overnight incubation (Supplementary Figure 2). NMR analysis of the hydrolysis mixture showed rapid disappearance of the signal at 4.04 ppm, indicating that the readily hydrolyzed methyl ester was in the phenazine portion. A similar behavior has been reported that allowed transformation of streptophenazine A into C^15^. The existence of a free carboxyl group was confirmed by both amidation with ethylediamine and esterification with MeOH (Supplementary Figure 3). MS^2^ fragmentation of these derivatives confirmed that modifications were introduced into the N-acetyl-cysteine moiety, indicating that the free carboxylic group is located on this moiety (Supplementary Figure 3).

However, the NMR experiment described above was not sufficient to fully resolve the structure of **1** as no correlations among the different spin systems were detected in the 2D TOCSY and HMBC experiments. Moreover, out of the 30 carbons from the calculated molecular formula of **1**, one was missing in the NMR spectra and a second one had a very low intensity. It has been reported that rigid environments can result in missing NMR hydrogen and carbon signals, which can be observed by acquiring the spectrum at higher temperature^16^. If sulfur was directly linked to one of the missing carbons, it might hamper molecular mobility and lead to missing NMR signals. Thus we re-analysed **1** dissolved in DMSO-d_6_ with spectra acquired at stepwise increased temperatures up to 77°C / 350K. Experiments at 350K allowed observation of the missing proton H-1’ at 5.62 ppm and of the corresponding carbon at 45.9 ppm. These values agree with a thioether substituent at C1’, the presence of which was confirmed by the NOESY correlation between H-2’’ and H1’. Finally, the now visible COSY correlations among protons H1’, 2’ and 3’, together with NOESY correlations between the aromatic proton H-7 and H1’ and 2’, established that the alkyl chain was in the same position as in known members of the streptophenazine family (supplementary NMR data). Overall, **1** turned out to be identical to streptophenazine F, except for the OH at C1’ being replaced by an *N*-acetyl-cysteine moiety linked through a thioether bond.

Regarding the stereochemistry, the configuration at C1’ could not be resolved. Indeed, it was not possible to assign the relative stereochemistry of 1’ and 2’ from the coupling constant, as described for similar compounds^17^ by simply measuring the values of J_3_ coupling H1’-H2’. At room temperature, the H-1’ signal is missing, whereas at 350K it becomes a broad doublet with a J_3_ coupling constant H1’-H2’of 10 Hz. This value might be compatible with the larger values observed for the *unlike* oxygen-containing stereoisomers, in comparison with a lower value of 6-8 Hz observed for the *like* stereochemistry. However, the presence of the sulfur atom might alter, or even reverse, these values. Moreover, the broadness of the H-1’ signal does not allow to exclude the presence of an overlapping diastereomeric signal with a lower J. Thus, while we can assume that the C2’ stereocenter is *R* as in previously reported streptophenazines, we are unable to draw conclusions about the stereochemistry at C1’ and whether **1** is chirally pure. As discussed below, the cysteine moiety can be assumed to be in the *L* configuration.

The structure of **2** was easily deduced from its MS^2^ fragmentation and NMR fingerprint from a sample purified from a 5L culture as described in Experimental Section. MS fragmentation data suggested a structure like that of **1** with an aliphatic chain shorter by a methylene unit (14 Da), thus representing a derivative of streptophenazine A in which an *N*-acetyl-cysteine linked through a thioether bond replaces the OH at C1’. The NMR fingerprint of **1** is superimposable to that of **2** (Table 1), except for the expected missing signals in the aliphatic chain, but featuring the same branched aliphatic chain (Supplementary NMR data).

In conclusion, **1** and **2** were established to be streptophenazines with an *N*-acetylated cysteine attached to C1’ of the alkyl chain via thioether bridge as shown on Figure 2. It should be noted that the nomenclature of streptophenazines suffers from some double booking, as identical names have been assigned multiple times to different metabolites^13, 17-21^ (Supplementary Table 2). For this reason, we prefer to name the compounds reported herein as *N*-acetyl-cysteinylated streptophenazines A and F.

**Figure 2.**
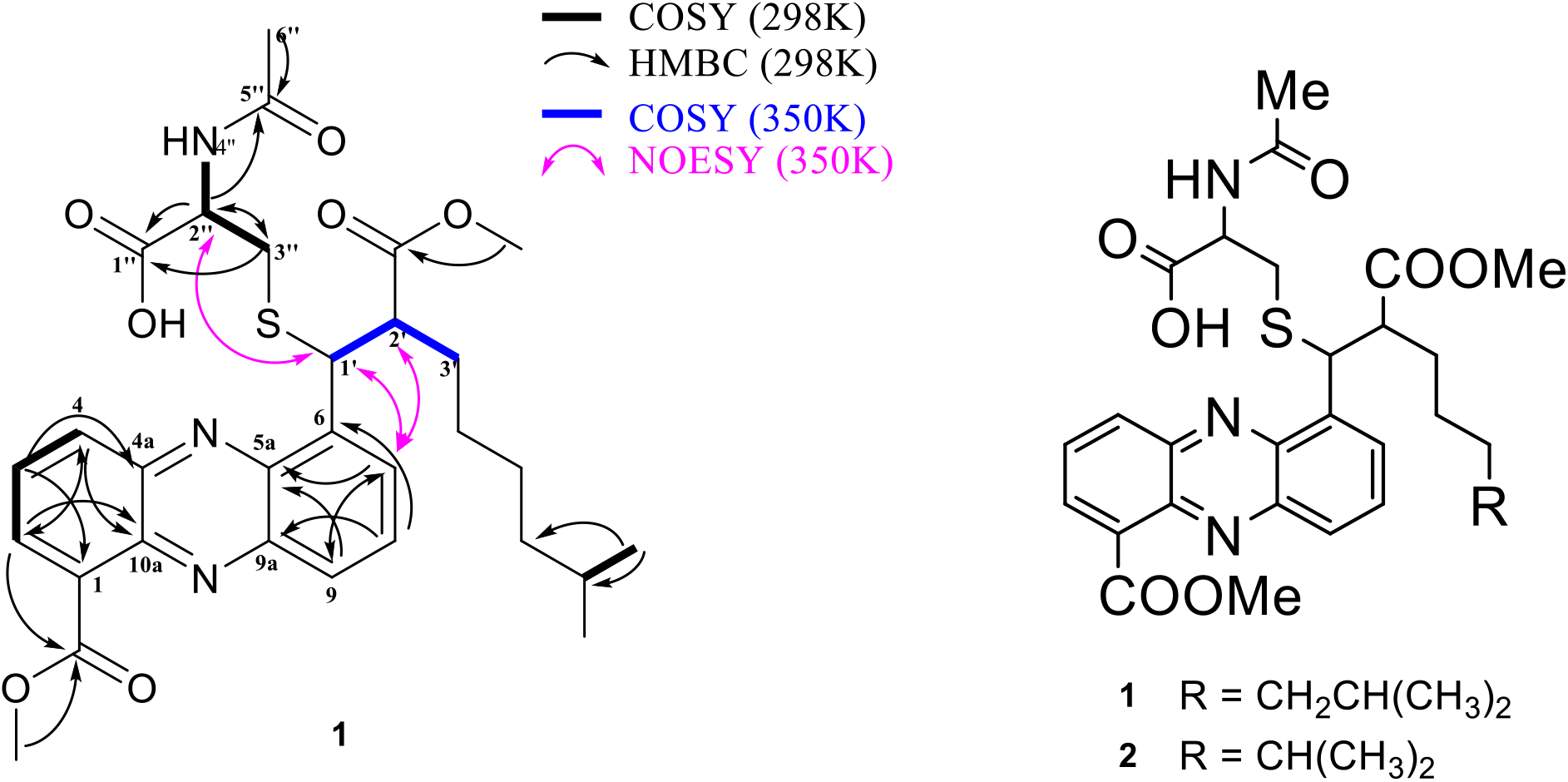
Left: selected NMR correlations for compound **1**. Right: structures of compounds **1** and **2.**

### Bioactivity and cytotoxicity

The aim of this study was to find novel antimicrobials using a bioactivity-independent approach. Indeed, compound **1** was found to suppress growth of *Streptococcus pneumoniae* and of *Staphylococcus aureus* with MICs of 12 and 43 μM, respectively (Figure 3A). Some activity was also observed against *Micrococcus luteus* (68 μM), *Escherichia coli ΔtolC* (291 μM), *Pseudomonas aeruginosa* and *Enterococcus faecium* (428 μM). The cytotoxicity of **1** was studied on CaCo-2 and HEK cells, which resulted in IC_50_’s of 154 and > 220 μM at 24h for CaCo-2 and HEK cells, respectively (Figure 3B).

**Figure 3.**
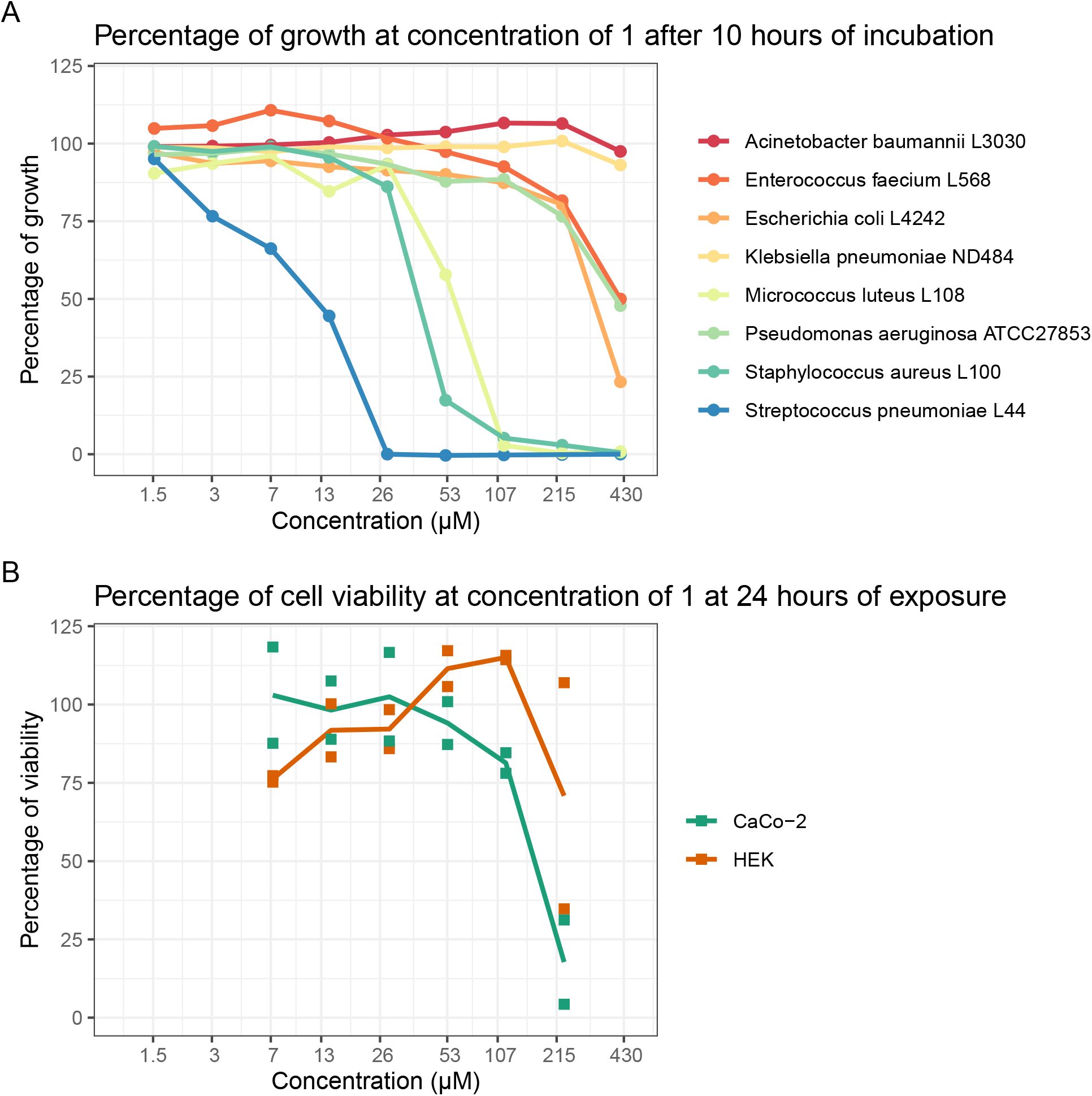
Antibacterial activity and cytotoxicity of compound **1**. Panel A: percentage of growth at 10h in the presence of **1**. *E. coli* L4242 is a *ΔtolC* derivative of strain M1522. Panel B: percentage of cell viability at 24 h in the presence of **1**. Note log scale on y axes.

### Biosynthetic gene cluster

Once **1** and **2** were known to belong to the streptophenazine class, we readily identified the BGC responsible for their formation from a draft genome of strain ID63040: antiSMASH^22^ version 6.0 equipped with MIBiG comparison tool^23^ proved to be very convenient for comparing the candidate BGC to those deposited in the MIBiG database, enabling us to identify BGCs which contain sets of similar genes.

Figure 4 presents a comparison between the BGC in strain ID63040 and two reference clusters: one for streptophenazines, from the marine *Streptomyces* sp. CNB-091^13^, and one for lomofungin, from *Streptomyces lomondensis* S015^24, 25^. The two streptophenazine BGCs are highly syntenic, sharing genes in similar order and orientation, apart from two insertions in the ID63040 BGC: a three-gene cassette, which is however present in the lomofungin BGC; and a regulator (yellow gene marked with an asterisk in Figure 4; see below). The three-gene cassette, which encodes for *S*-adenosylmethionine (SAM) synthetase, PfkB domain protein and methylenetetrahydrofolate reductase, is likely involved in the cofactor re-generation for the SAM-dependent methyltransferase. A methyltransferase is present in all the three BGCs of Figure 4 and is likely responsible for installing the methyl ester(s) on these molecules.

**Figure 4.**
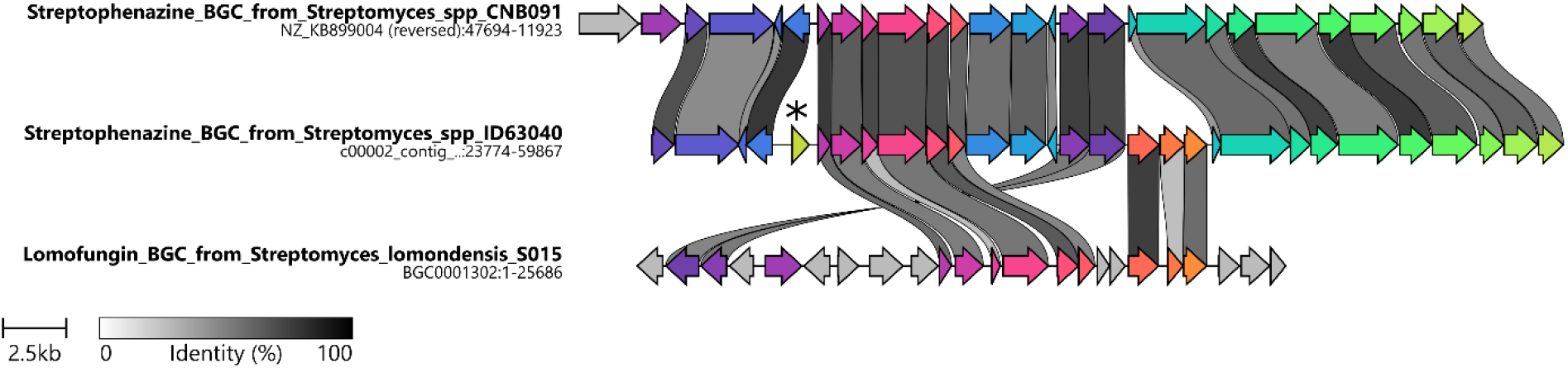
Comparison of streptophenazine BGC from strain ID63040 (middle) with reference streptophenazine BGC (top) from strain CNB091 and with lomofungin BGC from *Streptomyces lomondensis* S015 (bottom).

The regulatory gene embedded in the ID63040 BGC (locus tag ctg1_5) precedes the six-gene cassette responsible for synthesis of the phenazine core. While a homolog of this gene is present in the strain CNB-091, it is instead located four genes downstream from the left end of the CNB-091 streptophenazine BGC, in a closely linked BGC for the biosynthesis of mayamycin - a structurally unrelated aromatic polyketide (Supplementary Figure 4). Interestingly, two precedents exist of (strepto)phenazine BGCs prioritized for heterologous expression, yet production was achieved only if the clusters were refactored by inserting strong promoters in front of the phenazine gene cassette^13, 26^ – the exact same position where the regulatory gene naturally lies in the ID63040 BGC. This regulator shows 68% identity to JadR1, an activator of the jadomycin BGC and member of the OmpR family, referred to as “atypical response regulators”^27^. Interestingly, JadR1 has been shown to respond to late products in the jadomycin pathway^28^. Thus we consider ctg1_5 to be responsible for constitutive production of streptophenazines in strain ID63040.

We then wondered about the co-occurrence of ctg1_5 in streptophenazine BGCs in genomic sequences. Using the approach described in Experimental Section, we detected four BGCs containing both the phenazine gene cassette and the PKS portion, indicative of a streptophenazine BGC, out of a dataset of 561 *Streptomycetales* genomes. In only one case was a ctg1_5 homolog embedded in the streptophenazine BGC, while in three other cases it was associated with a closely linked mayamycin BGC (Supplementary Figure 4). Without associated metabolomics data, we can only speculate that the strain carrying ctg1_5 homolog within the streptophenazine BGC might not require refactoring for streptophenazine production.

During their extensive analysis of streptophenazine metabolites from the native host and after introducing a refactored BGC into *Streptomyces coelicolor* M1146, Bauman et al. observed streptophenazines A, F and G as main congeners^13^. Upon close inspection of their data we found that the strain carrying the refactored BGC also produced small amounts of compounds with *m/z* of 584.2413 and 570.2228 [M + H]^+^, which are consistent with **1** and **2**, respectively. Indeed, when the fragmentation patterns of the metabolites in both datasets were analyzed with the MASST tool^29^, both **1** and **2** were found in datasets associated with results published by Bauman et al.^13^ with a cosine score of 0.78 and 0.74 for *m/z* 570.1783 and 584.1971, respectively (Supplementary Table 3).

It should be noted that Bauman et al. characterized *N*-formyl-glycine adducts at C’1-OH of streptophenazines A and B as minor components of the complex and proposed that the ester ligase Spz15 is responsible for making this bond. Thus, position C1’ can undergo different modifications in different strains: esterification of the hydroxyl as reported by Bauman et al. and thioether formation in the present case. However, we doubt that the Spz15 homologue present in the ID63040 BGC is making the thioether bond observed in **1** and **2** (see below).

### Congener abundance dynamics

Since compounds **1** and **2** are structurally related to streptophenazines F and A, respectively, we investigated whether the latter compounds could also be detected. Indeed, by close inspection of extracts from strain ID63040, we observed *m/z* [M + H]^+^ values of 439 (F) and 425 (A), which showed similar fragmentation patterns to **1** and **2**, respectively and superimposable UV spectra (Supplementary Figure 1). Using the Moldiscovery workflow on GNPS, we confirmed the presence streptophenazines A and F, as well as congeners B, C, D, E, G, K, N and O (data not shown). Of note, none of these molecules clustered into the same molecular family as **1** and **2** in molecular networks calculated on GNPS.

To determine the relative abundances of the different streptophenazine congeners in the mycelium extracts, we used UV measurements. Unlike the intensities of the *m/z* peaks that may depend strongly on ionization efficiency, dimerization, charges per molecule and adduct formation, UV measurements enable reliable quantification of target compounds that share a common chromophore, i.e. the phenazine ring. In Figure 5, we report the peak heights at 252 nm associated with each *m/z* value. Compounds **1** and **2** were the prevalent congeners at 24 hours, when A and F are hardly detectable. Only at 72 hours congener F became comparable in concentration to **1** and **2**. Thus, during exponential growth *Streptomyces* sp. ID63040 produced **1** and **2** as the main components of the streptophenazine complex, while streptophenazines A and F became significant metabolites only when the strain enters stationary phase.

**Figure 5.**
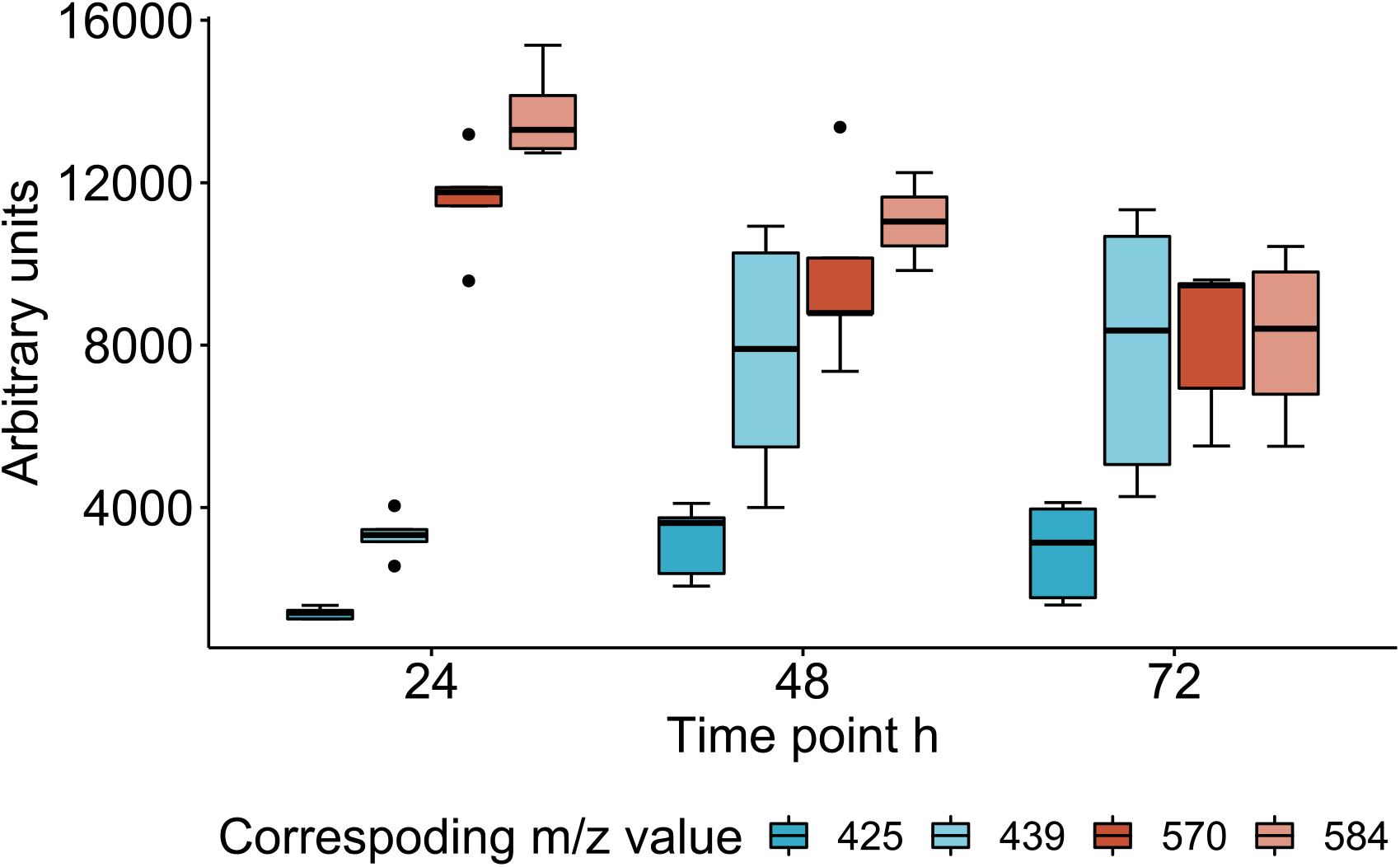
Abundances of streptophenazine congeners at 24, 48 and 72 h. *m/z* [M + H]^+^ 425, 439, 570 and 580 correspond to streptophenazine A, F, **1** and **2**, respectively. Results are expressed as average of five biological replicates run in parallel.

### On the possible role of streptophenazines

Phenazine is a moiety present in many known chemicals, both of synthetic and natural origin^30^. The properties of phenazine scaffold depend on pH, making it fit for a variety of tasks within and between microbial cells, ranging from carrying out redox functions and modulating gene expression to acting as signaling molecules and inhibiting growth of competitors^31^. Various mechanisms of action have been associated with phenazine antibiotics: e.g., myxin has been shown to intercalate with DNA, lomofungin inhibits RNA synthesis and pyocyanine provokes oxidative damage^32^. The best-known producers of naturally occurring phenazines are pseudomonads and streptomycetes^32^, the latter group featuring also their own special group of phenazine-derived metabolites named streptophenazines, which consist of an aliphatic tail added to the phenazine core by a polyketide synthase.

Previously, other phenazine natural products have been described carrying an *N*-acetyl-cysteine moiety attached through a thioether bridge to the aromatic core: examples are yorophenazine^33^, SB 212305^34^ and dermacozine J^35^ (Chart 1). Of note, all these molecules are devoid of the polyketide tail.

**Chart 1.**
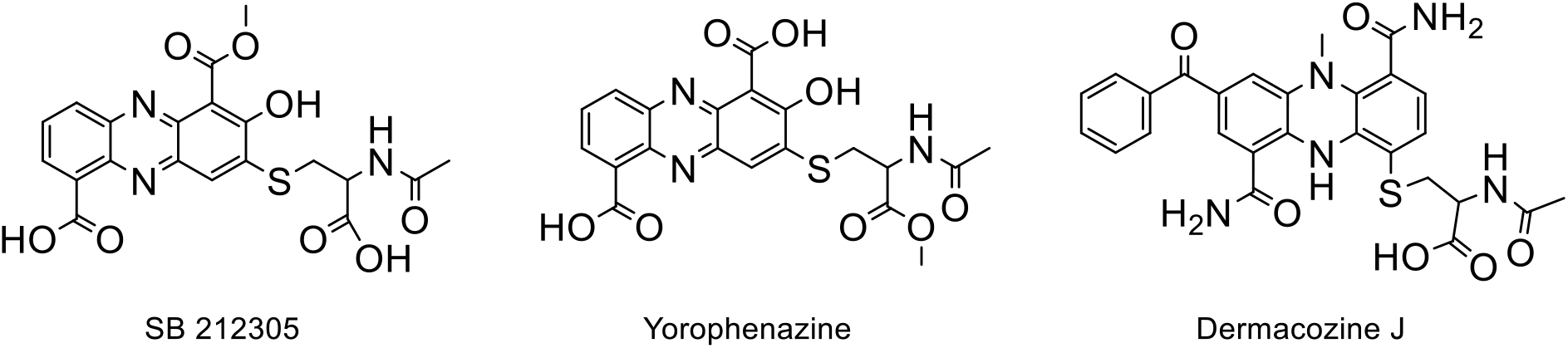
Structures of yorophenazine, SB212305 and dermacozine J^33–35^.

An *N*-acetylcysteine moiety has also been reported in several non-phenazine natural products^36–41^ and it has been hypothesized to result from mycothiol addition, a compound present in many actinobacteria where it is used to maintain cellular redox balance, to serve as a pool of stabilized cysteine and, in analogy to glutathione, to detoxify xenobiotics^42–44^. The mechanism of detoxification involves *S*-conjugation to the target molecule, followed by cleavage of the amide bond linking *N*-acetylcysteine to glucosamine in mycothiol. There are precedents for metabolites that have been shown or hypothesized to gain the *N*-acetylcysteine moiety from mycothiol: one such example is nanaomycin^45^, for which the authors described both the *N*-acetylcysteinylated and mycothiolated forms of the pyronaphtoquinone core. Thus, it is tempting to speculate that **1** and **2** are formed by such a mechanism.

While only few reports exist on the *in vivo* kinetics of *N*-acetylcysteine adduct(s) formation, this appears to be a slow process: for example, the mercapturic acid of granactin A appeared in cultures of *S. violaceruber* TÜ7 only after nine days, coinciding with disappearance of native granactin A^43^. By contrast, the cysteinylated products were the prevalent form during active growth of ID63040. A possible explanation of our findings might be that in strain ID63040 streptophenazines participate in an important redox function and, in doing so, are transformed into a toxic form that is readily neutralized by conjugation with mycothiol. As exponential growth slows or ceases, the role of streptophenazines in redox function decreases and congeners devoid of the *N*-acetyl-cysteine moiety (i.e., the true end product of the biosynthetic pathway) start to accumulate. If this hypothesis is correct, it would imply that streptophenazines in strain ID63040 play a role akin to primary metabolites, and hence their constitutive production through a BGC-embedded regulator. Of note, Price-Whelan et al.^31^ have previously stated that “phenazines blur the line between primary and secondary metabolism”.

## CONCLUSIONS

In summary, we have described two *N*-acetyl-cysteinylated streptophenazines with antibacterial activity identified through a metabolomics-first approach. Thus, it is possible to discover metabolites with antibacterial activity through a bioactivity-independent approach, even from a relatively small number of *Streptomyces* strains, a genus that has been intensively exploited for secondary metabolites, yet found to be by far the richest in terms of BGC diversity^47^. The prioritization criteria described herein represent just one such strategy, but we are confident that additional novel chemistry has been overlooked by our prioritization criteria, leaving room for further studies.

Our results also point out to the importance of the host in shaping the final metabolite profile. While it is well known that many BGCs are not expressed under laboratory conditions^46, 47^, BGC-specified metabolites can also undergo further spontaneous or enzymatic modifications as they are exposed to the cellular milieu^45, 48, 49^. The crosstalk between primary and secondary metabolism pathways has recently yielded new members even for one of the oldest known families of antibiotics^49^, underlining the somewhat arbitrary distinction between the two.

## EXPERIMENTAL SECTION

### General Experimental Procedures

LC-MS/MS analyses were performed on a Dionex UltiMate 3000 HPLC system coupled with an LCQ Fleet (Thermo Scientific) mass spectrometer equipped with an electrospray interface (ESI) and a tridimensional ion trap. The column was an Atlantis T3 C-18 5 μm X 4.6 mm X 50 mm maintained at 40 °C at a flow rate of 0.8 mL/min. Phase A was 0.05% trifluoroacetic acid (TFA), phase B was 100% acetonitrile. The elution was executed with a 14-min multistep program that consisted of 10, 10, 95, 95, 10 and 10% phase B at 0, 1, 7, 12, 12.5 and 14 min, respectively. UV-Vis signals (190-600 nm) were acquired using the diode array detector. The *m/z* range was 110-2000 and the ESI conditions were as follows: spray voltage of 3500 V, capillary temperature of 275 °C, sheath gas flow rate at 35 units and auxiliary gas flow rate at 15 units.

High resolution MS spectra were recorded at Unitech OMICs (University of Milano, Italy) using a Dionex Ultimate 3000 HPLC system coupled with a Triple TOF^®^ 6600 (Sciex) equipped with an ESI source. The experiments were carried using the same column and eluting system as described above. The ESI parameters were the following: curtain gas 25 units, ion spray voltage floating 5500 v, temperature 50 °C, ion source gas1 10 units, ion source gas2 0 units, declustering potential 80 v, syringe flow rate 10 μL/min, accumulation time 1 sec.

^1^H and ^13^C 1D and 2D NMR spectra (COSY, TOCSY, NOESY, HSQC, HMBC) were measured in DMSOd6 at the indicated temperature using a Bruker Advance II 300 MHz spectrometer.

### Production and purification of streptophenazines

For production of **1**, 1.5 ml of the −80°C stock culture was inoculated into 100 ml of AF medium (dextrose monohydrate 20 g L-1, yeast extract 2 g L-1, soybean meal 8 g L-1, NaCl 1 g L-1, and CaCO3 3 gL-1, pH 7.3 prior to autoclaving^4^) in two 300 ml baffled flasks and grown for three days at 30°C at 200 rpm. 30 ml of AF culture was transferred into five 2-L flasks with four baffles containing 500 ml M8 medium (dextrose monohydrate 10 g L-1, yeast extract 2 g L-1, meat extract 4 g L-1, soluble starch 20 g L-1, casein 4 g L-1, and CaCO3 3 g L-1, pH 7.2 prior to autoclaving^4^). The culture was harvested at 72 hours, filtered through Whatman paper and mycelium was extracted with 430 ml 100% EtOH by shaking for 1 hour at 30°C. The extract was filtered through Whatman paper and through 0.2 μm syringe filter and dried with a rotary evaporator.

For production of **2**, 0.75 ml of frozen culture was and inoculated into 50 mL baffled Erlenmeyer flasks containing 10 mL of medium AF. After 72h at 30°C on an orbital shaker at 200 rpm, 5 mL of the culture was transferred into each of six baffled 500-mL Erlenmeyer flasks containing 150 mL of fresh AF medium. After another 72 hours at 30 °C, 500 ml of culture was used to inoculate a 5L fermenter (New Brunswick, BioFlo^®^/CelliGen^®^ 115) containing 4.5 L of medium M8 supplemented with 0.5 ml/L polypropylene glycol (Sigma) as antifoaming agent. The fermenter was operated under following parameters: agitation 400 rpm, temperature 30°C, air flow 5 ml/min. The culture was harvested at 48 hours, filtered through Whatman paper and the resulting 1.2 L of mycelium was extracted with 1 L 100% EtOH by shaking overnight at room temperature. The extract was filtered three times through Whatman paper and dried with a rotary evaporator.

After drying, the residues were dissolved in 50% dimethylformamide (DMF) and fractionated separately using a TeledyneISCO CombiFlash MPLC mounted with Biotage SNAP Ultra C18 25um 60g column with detection at 252 nm. Phases A and B were 0.05% TFA and MeCN, respectively. The gradient used was 5%, 5%, 60%, 95% and 95% phase B at 0, 5, 10, 30 and 35 min, respectively. Throughout the purification procedure, the presence of **1** and **2** was monitored by LC-MS.

### Derivatization of 1 and 2

The number of free carboxyl groups in **1** and **2** was established by methylation and amidation reactions. Methylation: a 50-μl solution of compounds **1** and **2** in MeOH was treated with 2 μl H_2_SO_4_ at room temperature. Amidation: a dried mixture of compounds **1** and **2** was dissolved in 100 μl dry DMF and treated with 2 μl of ethylene diamine diluted 1:50 in dry DMF in the presence of few crystals of HATU.

The number of methyl esters was confirmed by basic hydrolysis. A dried sample of compounds **1** and **2** was treated with 2N NaOH at room temperature and the reaction mixture was analyzed by LC-MS prior to, and at 5 minutes, 1 hour and 24 hours after NaOH addition.

### Bioassays

The antibacterial activity of compound **1** was determined against strains from the NAICONS collection of bacterial pathogens as follows: 90 μl of a 1×10^5^ CFU/ml bacterial suspension in the appropriate medium was dispensed into each well of a 96-well plate containing 10 μl of compound **1**, dissolved at 428 μM in 2.5% DMSO and serially diluted 1:2 with 2.5% DMSO. The medium was cation-adjusted MHB for all strains except for *S. pneumoniae*, for which THB was used. Plates were incubated in a Synergy 2 (Biotek) microplate reader with readings at 595 nm registered every hour.

For cytotoxicity assays, HEK 293 cells and CaCo-2 cells were cultured in DMEM/F-12 and DMEM (Gibco), respectively, supplemented with 10% fetal calf serum (FCS) and 1% of Penicillin-Streptomycin mixture (Gibco) at 37°C 5% CO_2_. Cells were seeded in 96-well plates at a concentration of 1 x 10^5^ cells per well, the outer wells were filled with PBS to avoid disturbances due to evaporation. After 24 hours incubation, cells were confluent. Two-fold dilutions of the compounds were made in a range of 214-6.7 μM, in exposure medium. The exposure medium consisted of DMEM without phenol red (Gibco), without supplementation with FCS or antibiotics.

The growth medium from the plates was aspirated and replaced with 100 μl exposure medium containing different concentrations of the compounds. Three wells were used as a medium control, three as vehicle control (1% DMSO) and three as positive control for toxicity (20% DMSO). After 24 hours exposure, cytotoxicity was measured using Alamar Blue assay (Thermo Fisher Scientific) by adding 10μl of reagent per well, incubating 1 hour at 37ºC 5% CO_2_ and measuring fluorescence (λ_EX_ = 540 nm, λ_EM_ = 590 nm) on a SpectraMax M5 (Molecular Devices). The IC_50_ was calculated using GraphPhad Prism 9 using a non-linear regression analysis: log(inhibitor) vs. response – variable slope (four parameters) without constraints.

### Bioinformatic analyses

From a draft genome sequence of *Streptomyces* strain ID63040, obtained using both Illumina short read sequencing and Oxford Nanopore Technologies (ONT) long read sequencing and consisting of 11 contigs with total length of 8,915,670 bp (K.V. and V.W, unpublished), BGCs were identified using the antiSMASH version 6.0 online tool^23^.To search for the occurrence of homologs of the regulator ctg1_5 in streptophenazine BGCs, we searched the antiSMASH-DB^28^ with KnownClusterBlast for BGCs matching both the phenazine and the PKS portion of the streptophenazine BGC BGC0002010. BGC alignments were visualized with Clinker^50^. The streptophenazine BGC from Streptomyces strain ID63040 was deposited to GenBank with accession No. OL619055.

The 16S rRNA gene sequence (GenBank accession No. OL423644) was determined and analyzed as described by Monciardini et al. 2002^51^.

## Supporting information

Supplemental NMR data

## ASSOCIATED CONTENT

### Supporting Information

Supplementary figures and tables (PDF)

Supplementary NMR data (PDF)

### Author Contributions

K.V. and S.D. conceived and designed the experiments; K.V., S.M., C.B., B.F.C. and V.W carried out the experimental work; K.V., M.S., M.S., S.M. and W.V. analyzed the data; K.V, S.M and S.D. wrote the paper. The manuscript was written through contributions of all authors. All authors have given approval to the final version of the manuscript.

### Notes

The authors declare no competing financial interest.

## ACKNOWLEDGMENTS

This work received funding from the European Union’s Horizon 2020 research and innovation programme under the Marie Sklodowska-Curie grant agreement No 765147. We thank Aigars Jirgensons and Hans-Georg Sahl for their contributions in improving the manuscript. KV thanks Simen Fredriksen for help with data handling.

## ABBREVIATIONS

BGC: biosynthetic gene cluster
NMFL: NAICONS metabolic fingerprint library
MPLC: medium pressure liquid chromatography

## Notes

### Competing Interest Statement

The authors have declared no competing interest.

