## Supplemental figures and tables for "*N*-acetyl-cysteinylated streptophenazines from *Streptomyces*"

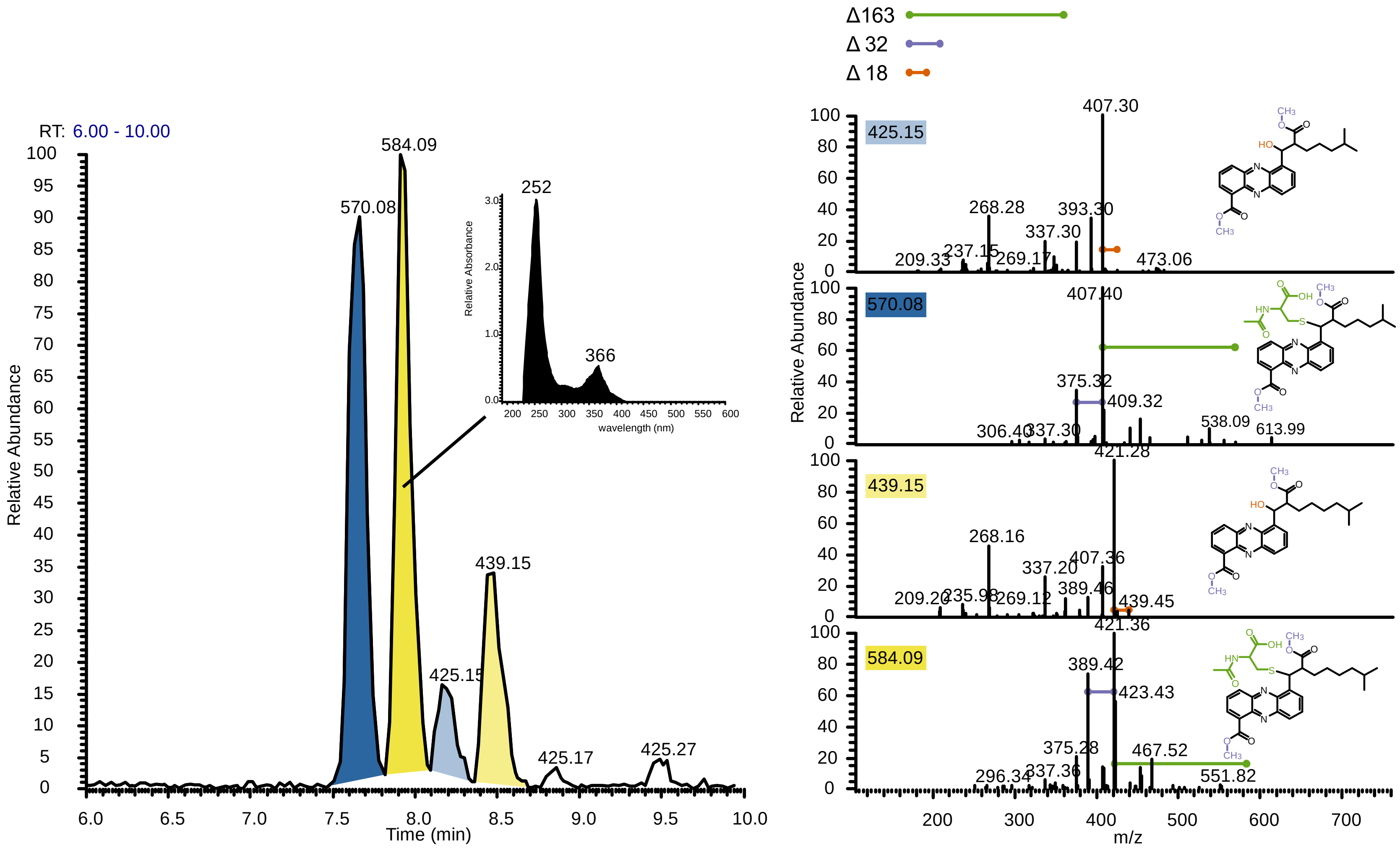


*Supplementary Figure 1. Fragmentations of **1** and **2** and their putative precursors. Left: EIC of m/z $[M + H]^+$ values 584.4, 570.4, 425.2 and 439.2., corresponding to **1** (yellow background) and **2** (blue background) or to their precursors (background in lighter shade). Embedded: UV-vis absorption spectrum corresponding to m/z 584.4 $[M + H]^+$. Right: fragmentation patterns of **1**, **2** and their precursors. Horizontal bars indicate the size of neutral losses with corresponding part highlighted in structure with same color. Data was acquired with low resolution LC-MS instrument.*

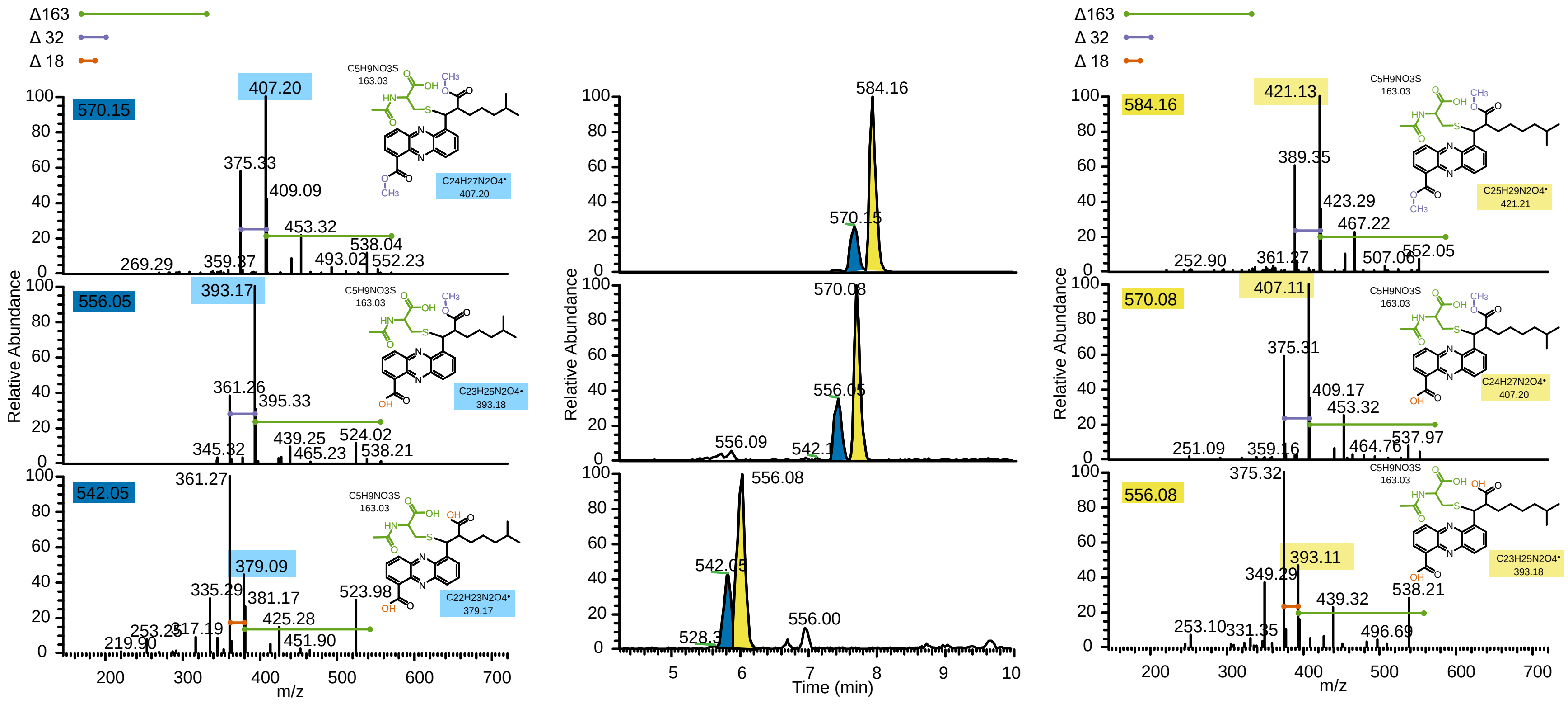


*Supplementary Figure 2. Alkaline hydrolysis of **1** and **2**. Middle panel: Extracted Ion Chromatogram (EIC) of m/z [M + H]+ values 584.4, 570.4, 556.4, 542.4 and 528.4., corresponding to **1** (yellow background) and **2** (blue background) or their derivatives. From top to bottom: analyses before treatment, 5 minutes and 24 hours after treatment with 2N NaOH. Left panel: fragmentation patterns of **2** and its derivatives. Right panel: fragmentation patterns of **1** and its derivatives. The bars reflect the size of neutral loss on left and right panels: green corresponds to a neutral loss of 163 (N-acetylcysteine), purple to 32 (methyl ester) and orange to 18 (water). Data was acquired with low resolution LC-MS instrument. Note that the different hydrolytic behavior of methyl esters (green ink) was established by an NMR analysis (see text).*

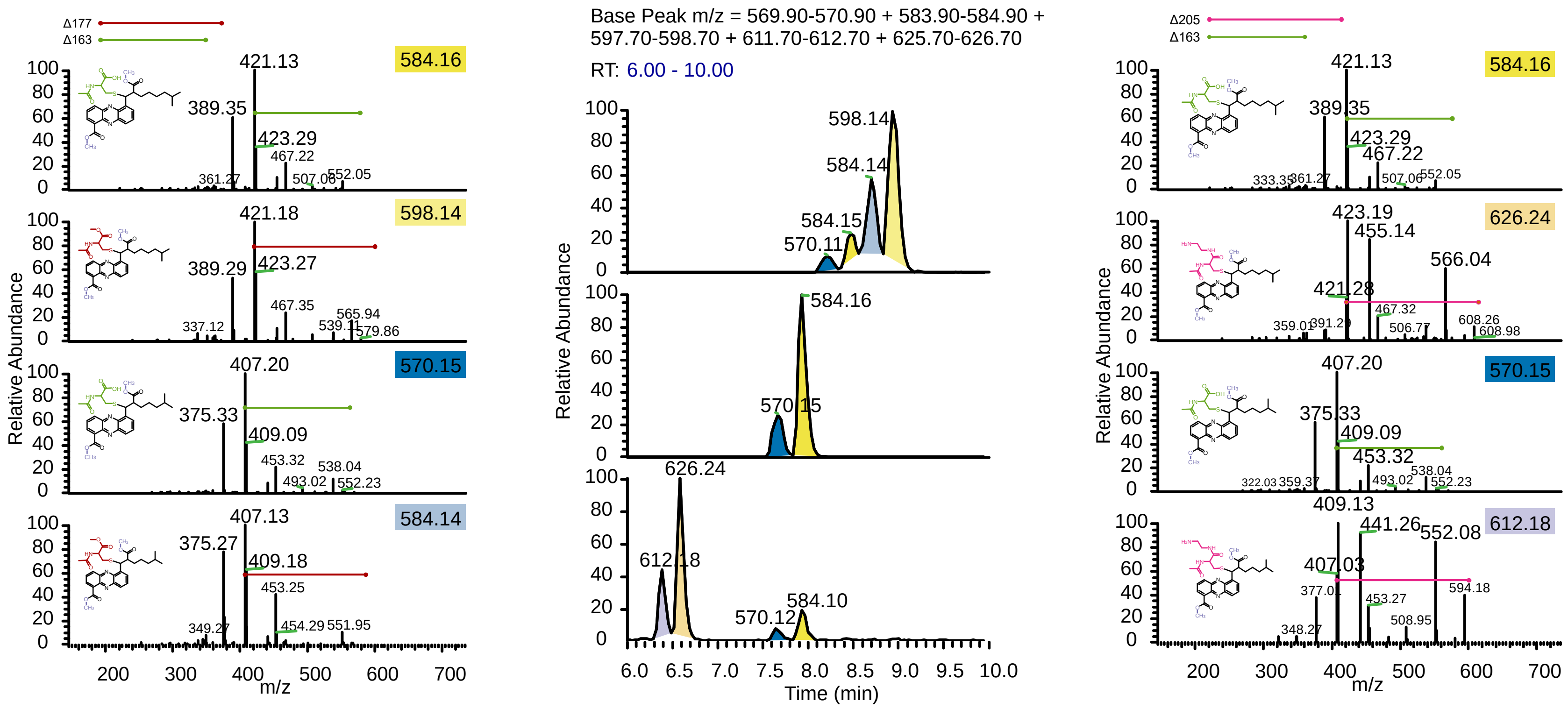


*Supplementary Figure 3. Derivatization of **1** and **2**. Middle: EIC of m/z [M + H]+ values 584.4, 570.4, 584.4, 598.2, 621.4 and 626.2, corresponding to **1** (yellow background) and **2** (blue background) or their derivatives. Left: MSMS patterns of native and methylated products. Right: MSMS patterns of native and amidated products. The bars reflect the size of neutral loss on left and right panels: green corresponds to a neutral loss of 163 (N-acetylcysteine), dark red to 177 (N-acetylcysteine + methyl) and pink to 205 (N-acetylcysteine + ethylenediamine). Data was acquired with low resolution LC-MS instrument.*

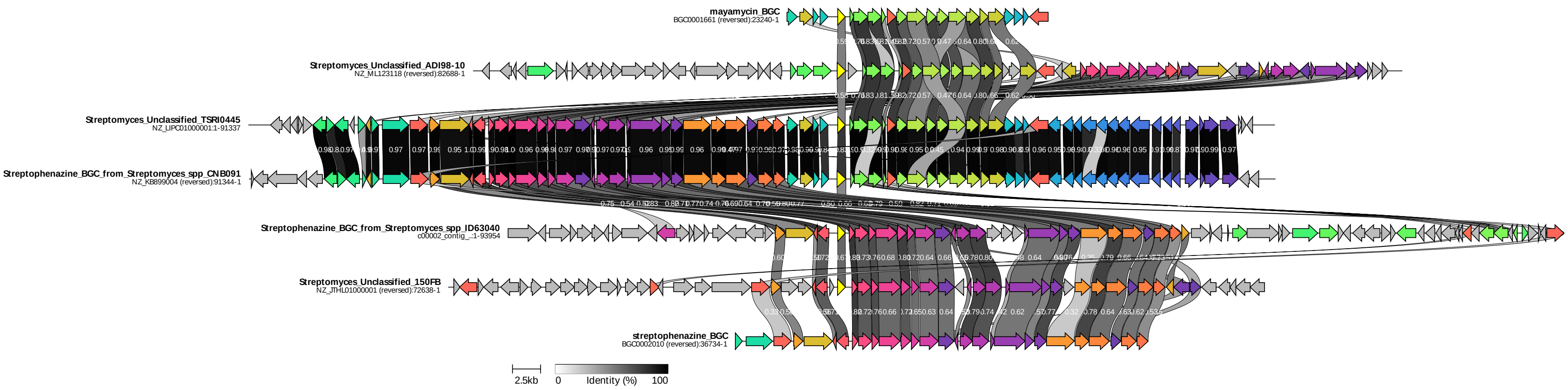


*Supplementary Figure 4. Comparison of streptophenazine BGC regions of strain ID63040 and Streptomyces streptophenazine BGCs from the antiSMASH database. The BGCs are anchored to the regulator ctg1_5 (yellow, marked with an asterisk) embedded in the ID63040 BGC. The mayamycin (top) BGC is also shown. Figure was generated with Clinker. White numbers on bands indicate identity of two genes.*

*Supplementary Table 1. Calculated molecular formulae for parent mass, selected fragments and neutral losses of compounds* ***1*** *(top) and* ***2*** *(bottom). Ppm values >20 are highlighted in red. Experiments were conducted with high resolution instrument.*

| m/z value | Neutral loss | C | H | N | O | S | Molecular formula | ppm | DBE |
|---|---|---|---|---|---|---|---|---|---|
| **584.2430** | | 30 | 38 | 3 | 7 | 1 | C30H38N3O7S | 4.9 | 13.5 |
| | 32.0235 | 1 | 4 | | 1 | | CH4O | -84.77 | 0 |
| 552.2195 | | 29 | 34 | 3 | 6 | 1 | C29H34N3O6S | 4.83 | 14.5 |
| | 131.0036 | 4 | 5 | 1 | 2 | 1 | C4H5NO2S | -3.81 | 3 |
| | 163.0271 | 5 | 9 | 1 | 3 | 1 | C5H9NO3S | -19.72 | 2 |
| 421.2159 | | 25 | 29 | 2 | 4 | | C25H29N2O4 | -6.85 | 12.5 |
| | 32.0268 | 1 | 4 | | 1 | | CH4O | 18.27 | 0 |
| 389.1891 | | 24 | 25 | 2 | 3 | | C24H25N2O3 | 6.64 | 13.5 |
| | 27.9950 | 1 | | | 1 | | CO | 3 | 2 |
| 361.1941 | | 23 | 25 | 2 | 2 | | C23H25N2O2 | 6.91 | 12.5 |
| | 110.1111 | 8 | 14 | | | | C8H14 | 14 | 2 |
| 251.0830 | | 15 | 11 | 2 | 2 | | C15H11N2O2 | 3.77 | 11.5 |

| m/z value | Neutral loss | C | H | N | O | S | Molecular formula | ppm | DBE |
|---|---|---|---|---|---|---|---|---|---|
| **570.2310** | | 29 | 36 | 3 | 7 | 1 | C29H36N3O7S | 6.32 | 13.5 |
| | 32.0275 | 1 | 4 | | 1 | | CH4O | 40.13 | 0 |
| 538.2035 | | 28 | 32 | 3 | 6 | 1 | C28H32N3O6S | 4.31 | 14.5 |
| | 131.0042 | 4 | 5 | 1 | 2 | 1 | C4H5NO2S | 0.77 | 3 |
| | 163.0317 | 5 | 9 | 1 | 3 | 1 | C5H9NO3S | 8.5 | 2 |
| 407.1993 | | 24 | 27 | 2 | 4 | | C24H27N2O4 | 5.45 | 12.5 |
| | 32.0268 | 1 | 4 | | 1 | | CH4O | 18.27 | 0 |
| 375.1725 | | 23 | 23 | 2 | 3 | | C23H23N2O2 | 4.35 | 13.5 |
| | 27.9947 | 1 | | | 1 | | CO | -7.67 | 2 |
| 347.1778 | | 22 | 23 | 2 | 2 | | C22H23N2O2 | 5.32 | 12.5 |
| | 96.0952 | 7 | 12 | | | | C7H12 | 13.52 | 2 |
| 251.0826 | | 15 | 11 | 2 | 2 | | C15H11N2O2 | 2.18 | 11.5 |

*Supplementary Table 2. Strains reported to produce streptophenazines and BGC sequence availability*

| Streptophenazines | Authors | Producer strain | Habitat | Sampling site | Host | BGC sequence availability |
|---|---|---|---|---|---|---|
| Streptophenazines A-H | Mitova, Lang et al. 2008 | Streptomyces sp. strain HB202 | marine | Germany, Baltic Sea | marine sponge Halichondria panicea | No |
| Revised structure of Streptophenazine A | Yang, Jin et al. 2011 | Streptomyces sp. strain HB202 | marine | Germany, Baltic Sea | marine sponge Halichondria panicea | No |
| Streptophenazines I-L | Bunbamrung, Dramae et al. 2014 | Streptomyces sp. BCC21835 | terrestrial | Thailand, Khao Khitchakut National Park, Chanthaburi province | soil | No |
| Streptophenazines I-K | Kunz, Labes et al.2014 | Streptomyces strain HB202 | marine | Germany, Baltic Sea | marine sponge Halichondria panicea | No |
| Streptophenazines M-O | Liang, Chen et al. 2017 | Streptomyces sp. 182SMLY | marine | China, East China Sea | sediment | No |
| Streptophenazines P-R | Bauman, Li et al. 2019 | Streptomyces sp. CNB-091 | marine | USA, Florida Keys | jellyfish Cassiopeia xamachana | Yes |

*Supplementary Table 3. Hits for N-acetyl-cysteinylated streptophenazines **1** and **2** in public databases.*

| m/z | MASST dataset ID | Source | Matched peaks | Cosine score |
|---|---|---|---|---|
| 570.1783 | MSV000083082:M1146-pKDB03.mzML | Streptomyces (NCBITaxon:1883) | 14 | 0.78 |
| 584.1971 | MSV000083082:M1146-pKDB03.mzML | Streptomyces (NCBITaxon:1883) | 10 | 0.74 |
