## Supplemental NMR data for "*N*-acetyl-cysteinylated streptophenazines from *Streptomyces*"

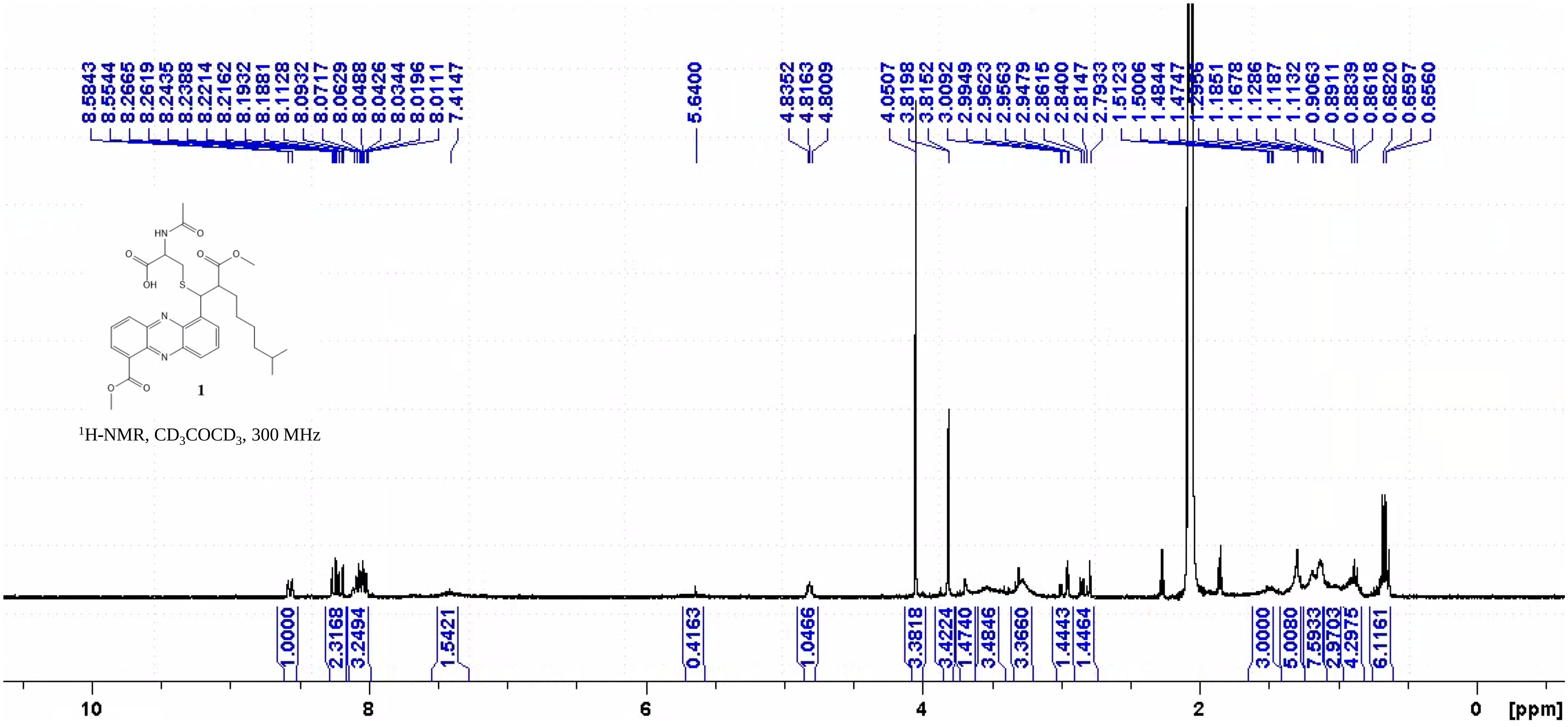


Supplementary Figure I: $^1$H-NMR of **1** in acetone-$d_6$ at 300K

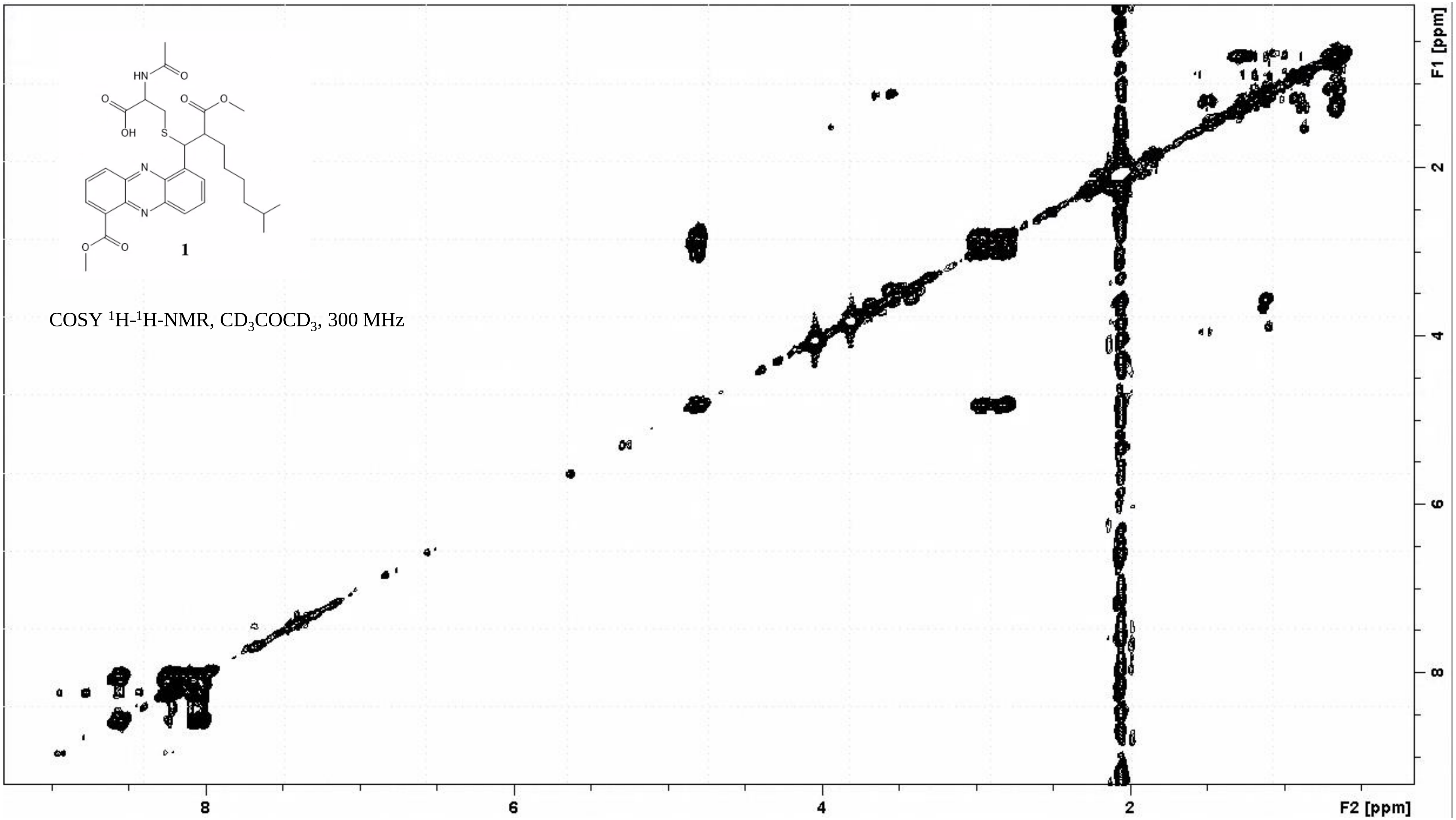


Supplementary Figure II: COSY of **1** in acetone-$d_6$ at 300K

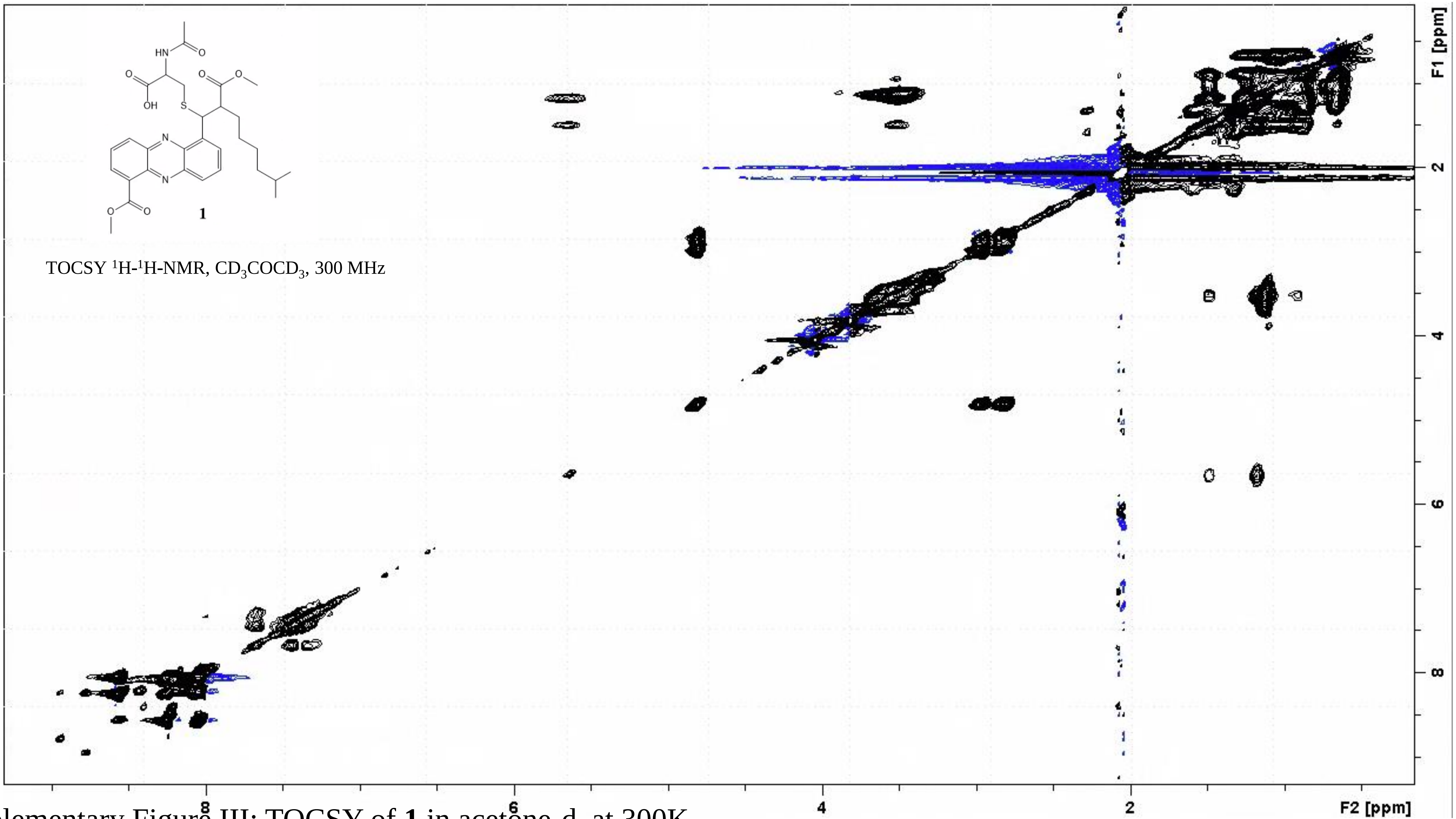


Supplementary Figure III: TOCSY of **1** in acetone-d$_6$ at 300K

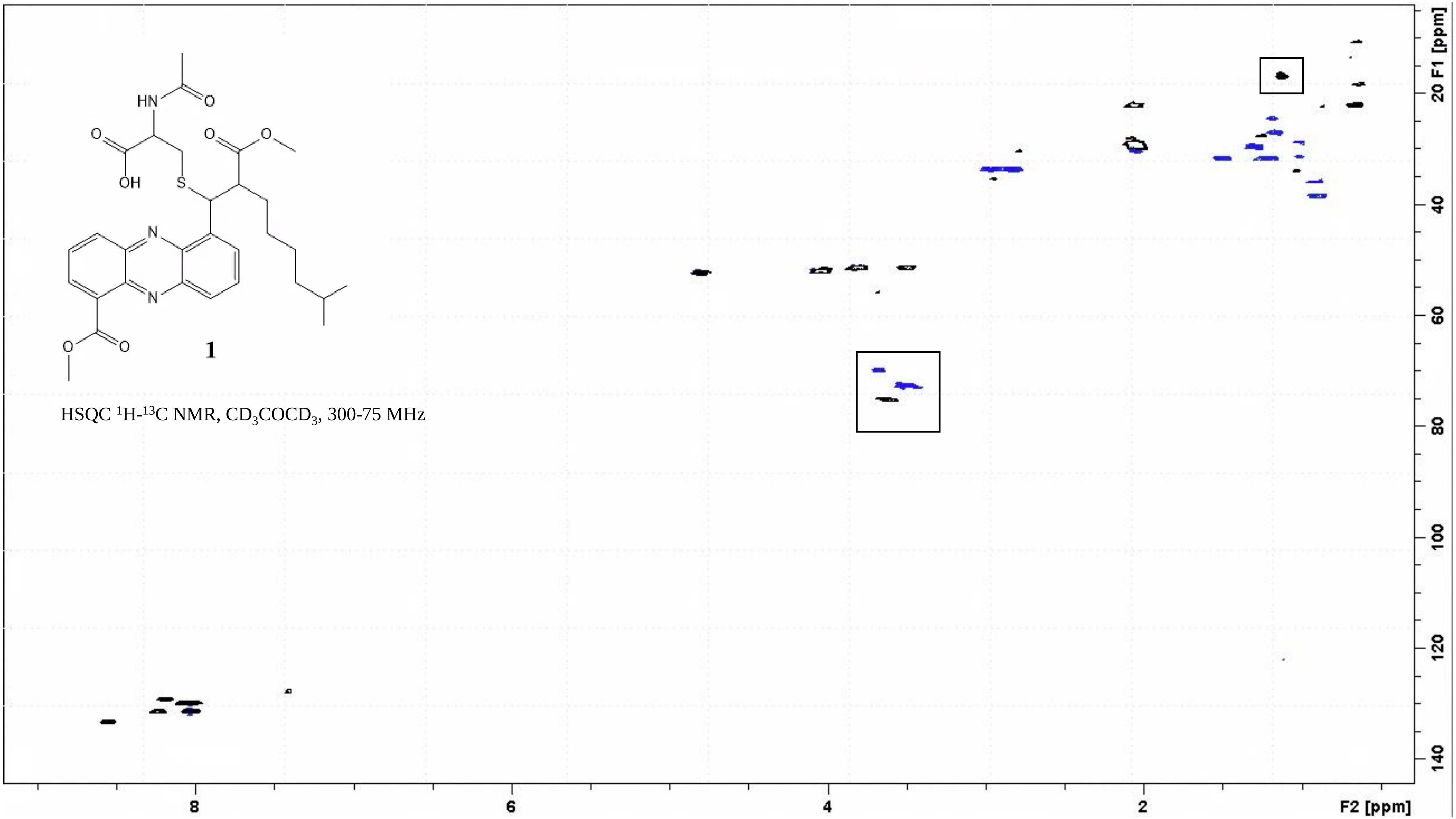


Supplementary Figure IV: HSQC of **1** in acetone-$d_6$ at 300K, in the boxes PPG impurities signals

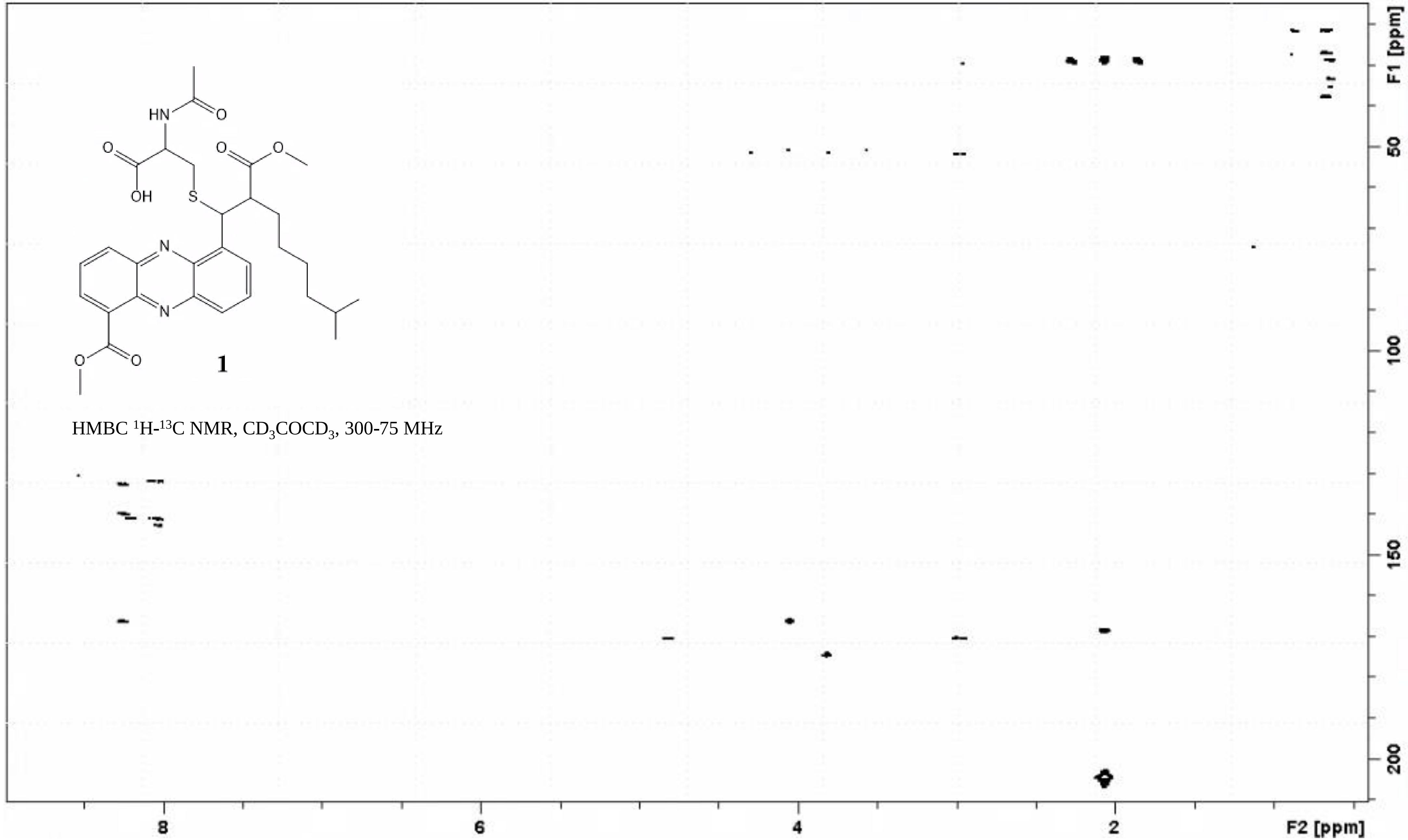


Supplementary Figure V: HMBC of **1** in acetone-$d_6$ at 300K

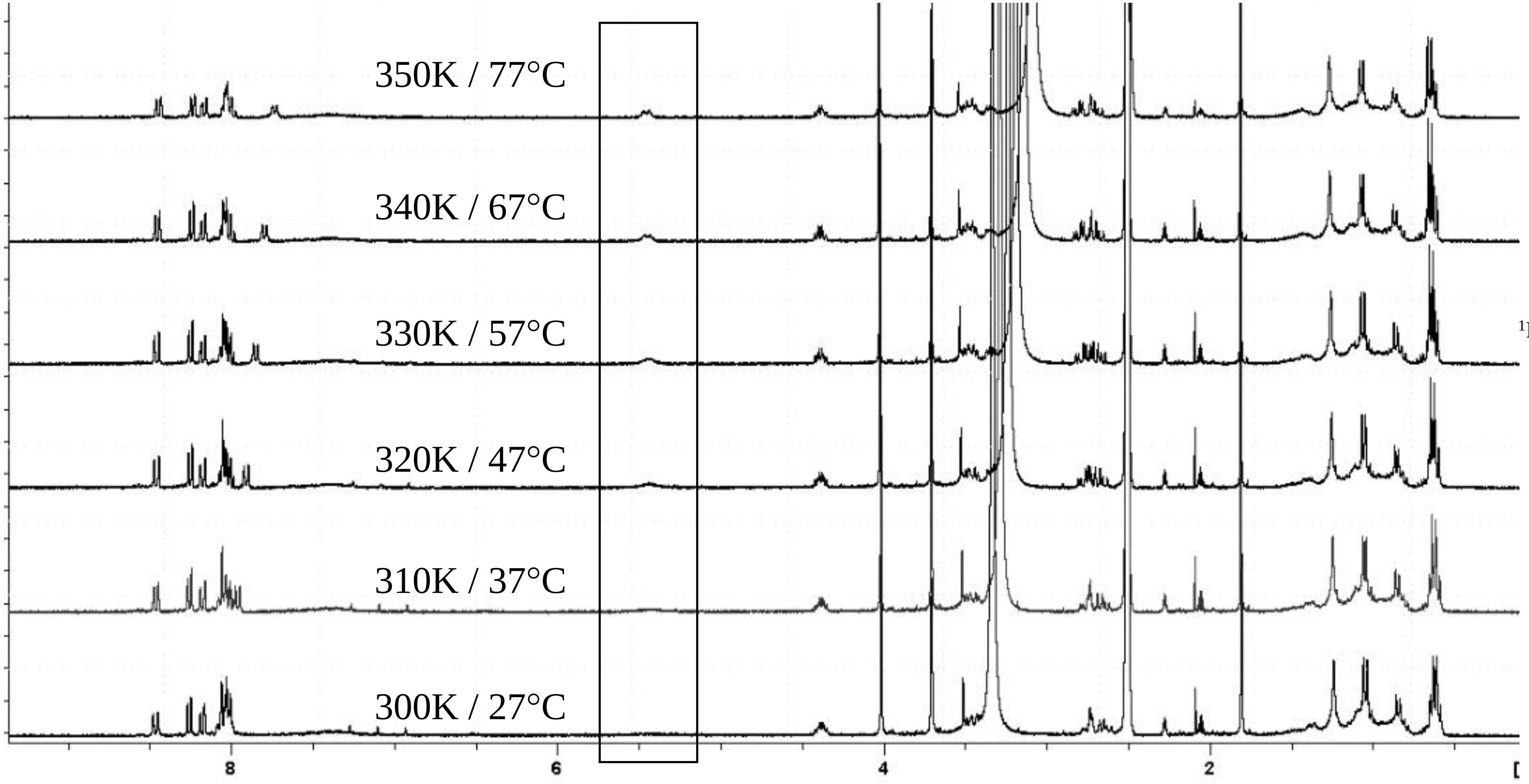


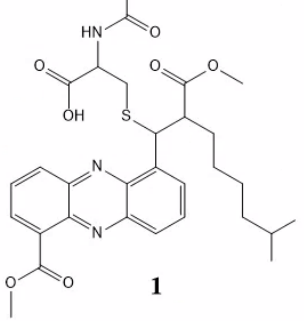


**1**

$^{1}$H-NMR, $CD_3SOCD_3$, 300 MHz

Supplementary Figure VI: $^{1}$H-NMR of **1** in dmso-$d_6$ while increasing temperature from 300K to 350K

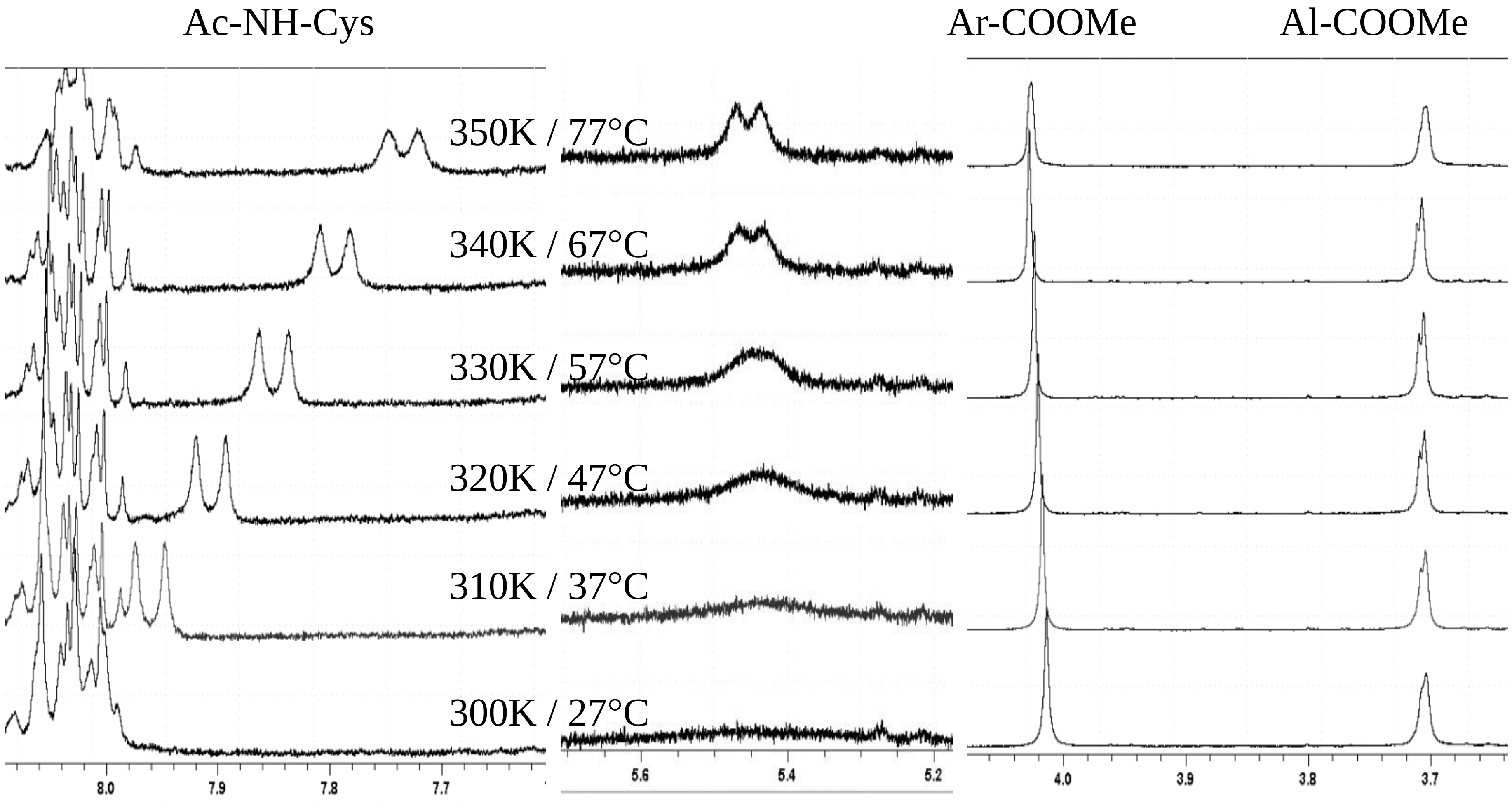


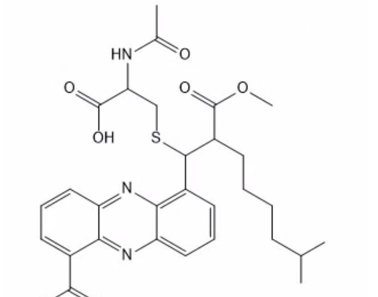


**1**

$^1$H-NMR, $CD_3SOCD_3$, 300 MHz

Supplementary Figure VII: $^1$H-NMR of **1** in dmso-$d_6$ while increasing temperature from 300K to 350K: selected signals

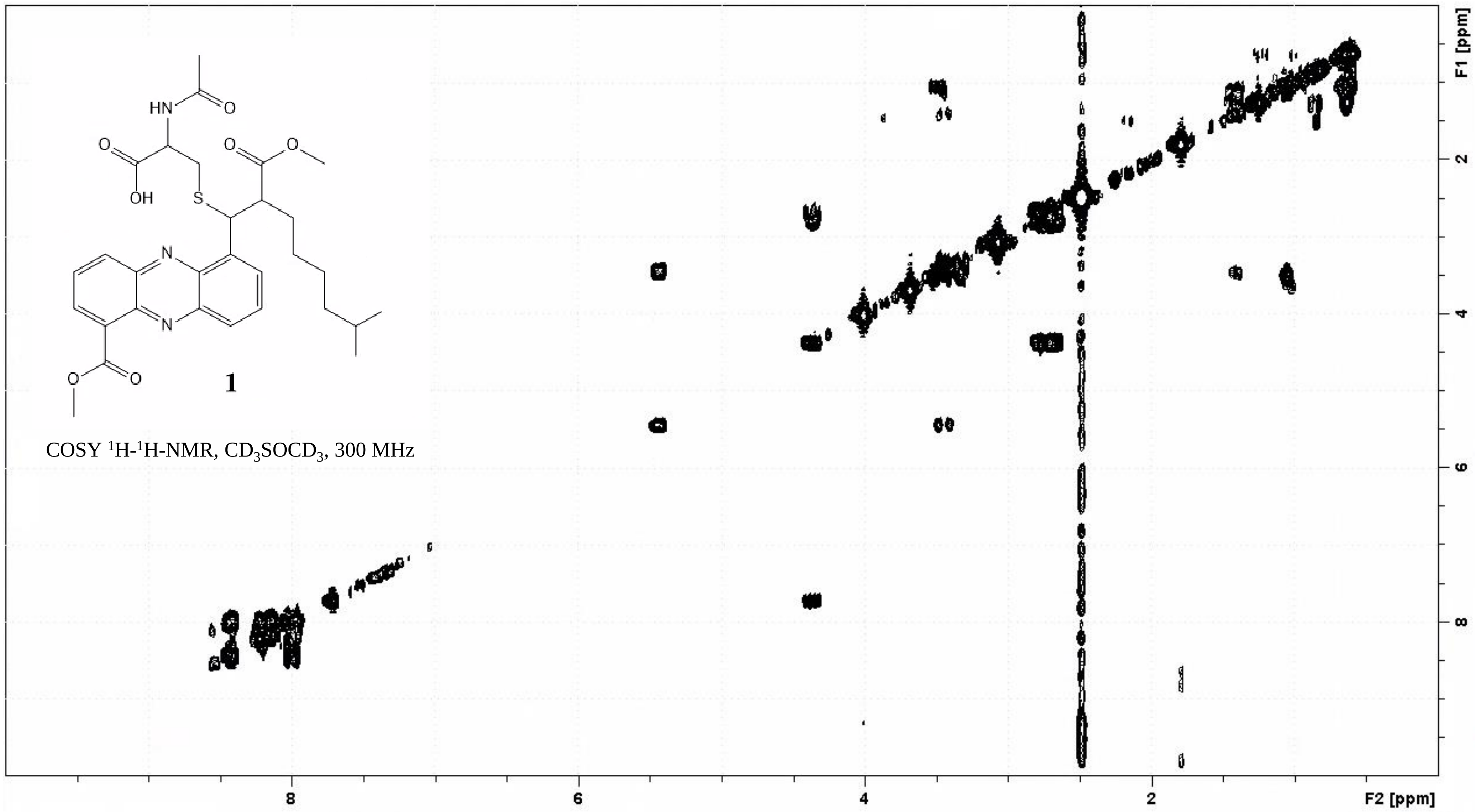


Supplementary Figure VIII: COSY of **1** in dmso-$d_6$ at 350K / 77°C

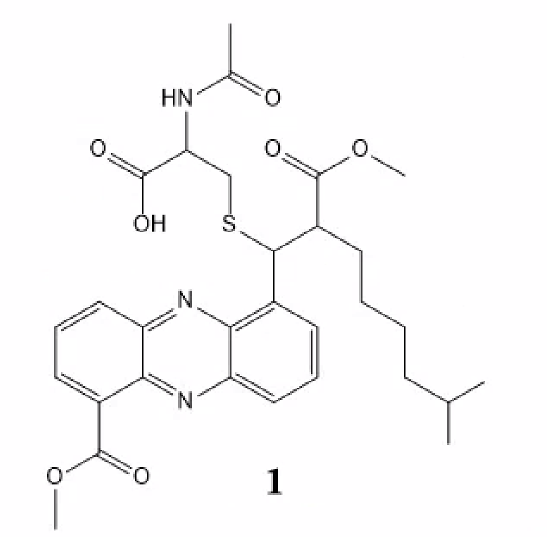


**1**

TOCSY $^1H$-$^1H$-NMR, $CD_3SOCD_3$, 300 MHz

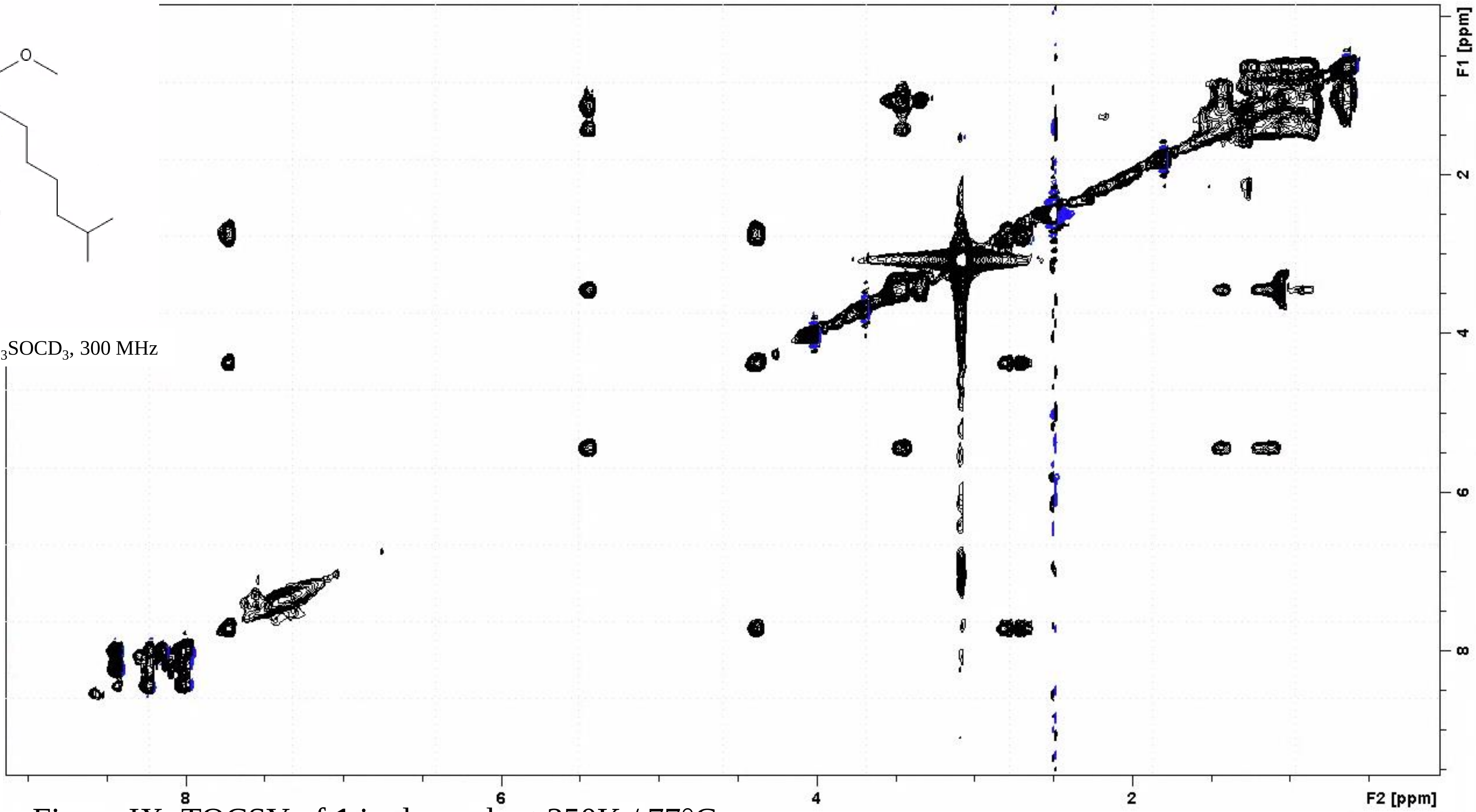


Supplementary Figure IX: TOCSY of **1** in dmso-$d_6$ at 350K / 77°C

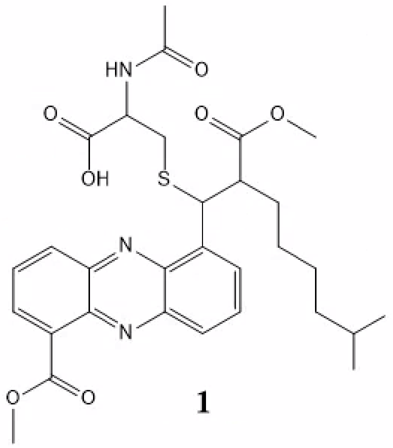


**1**

NOESY $^{1}H$-$^{1}H$-NMR, $CD_3SOCD_3$, 300 MHz

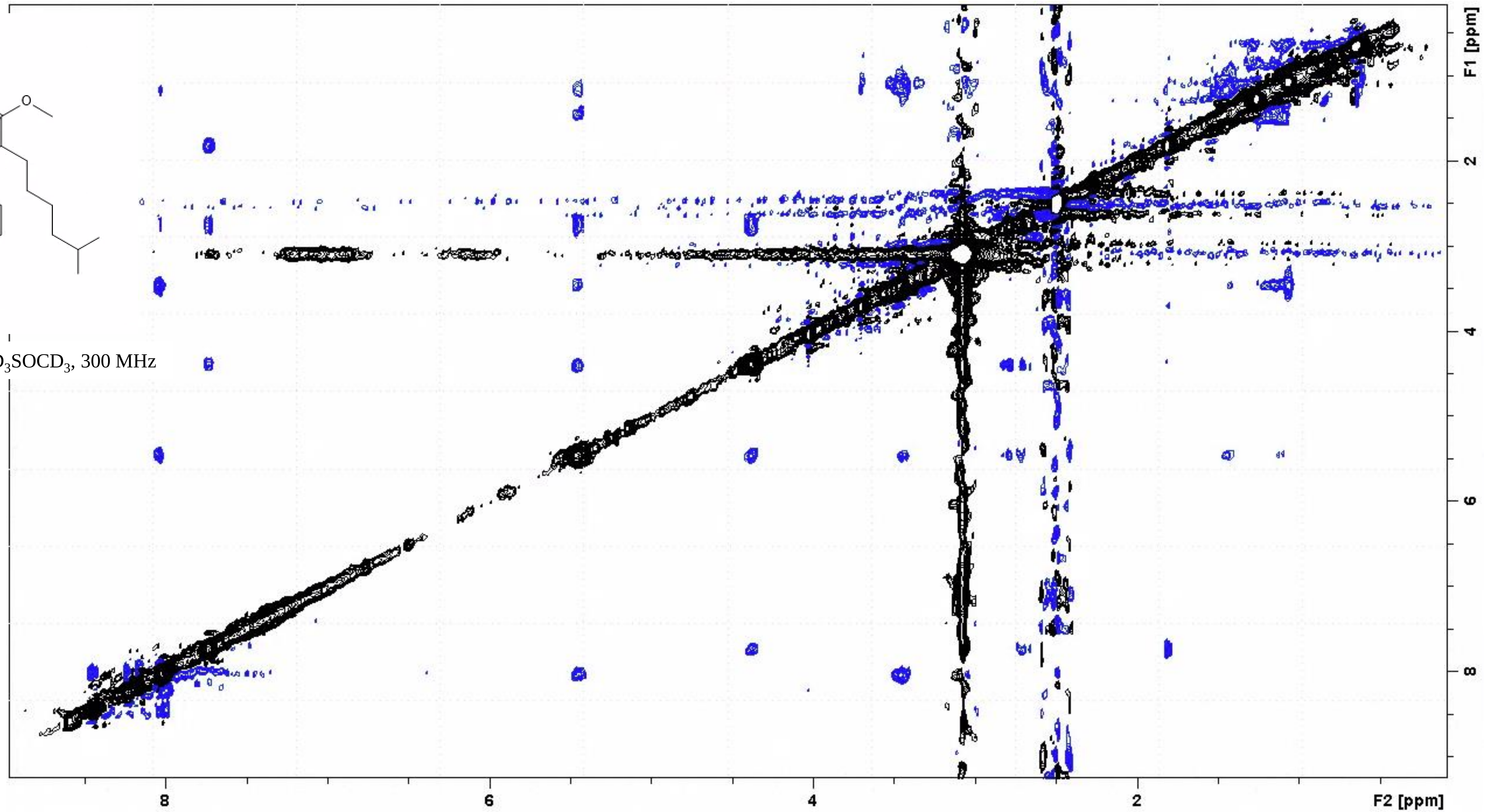


Supplementary Figure X: NOESY of **1** in dmso-$d_6$ at 350K / 77°C

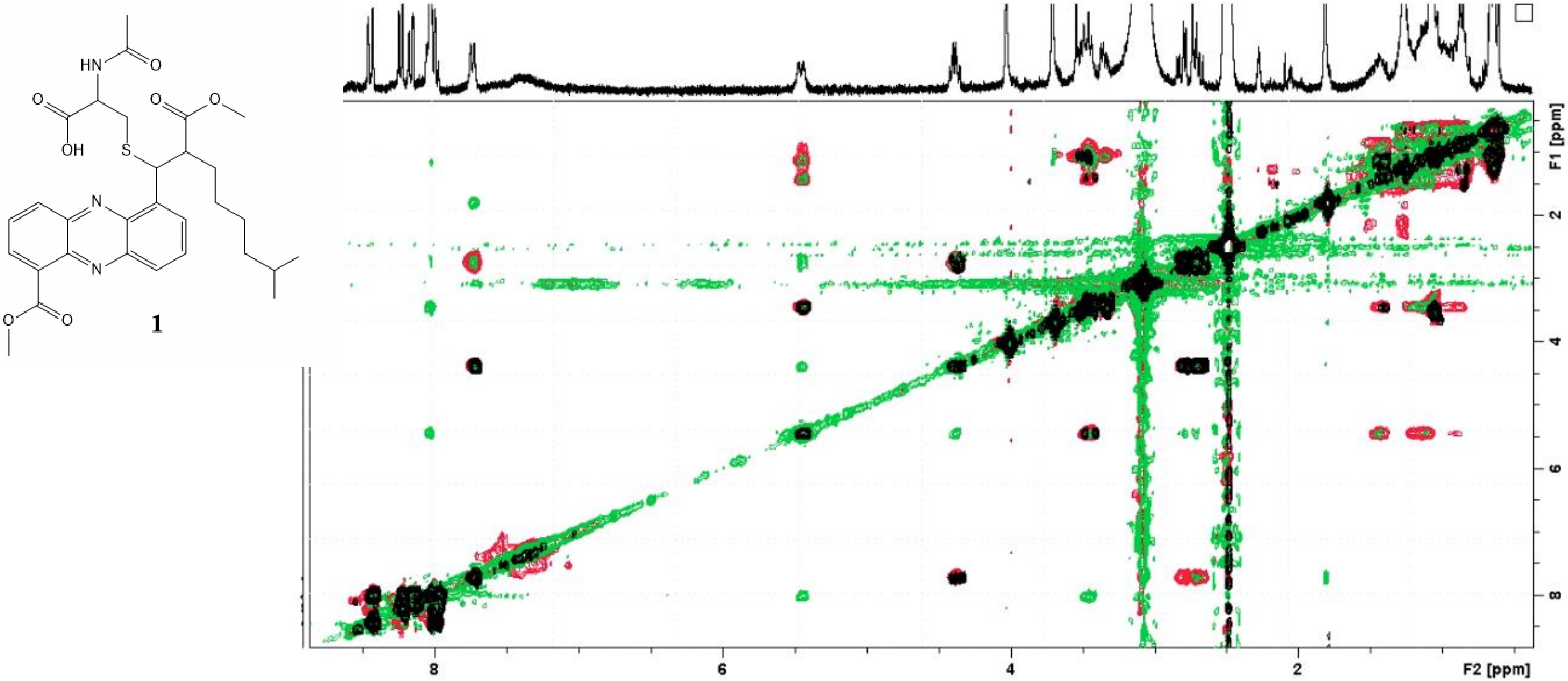


1

Supplementary Figure XI: COSY (black), TOCSY (red) and NOESY (green) of **1** in dmso-$d_6$ at 350K / 77°C

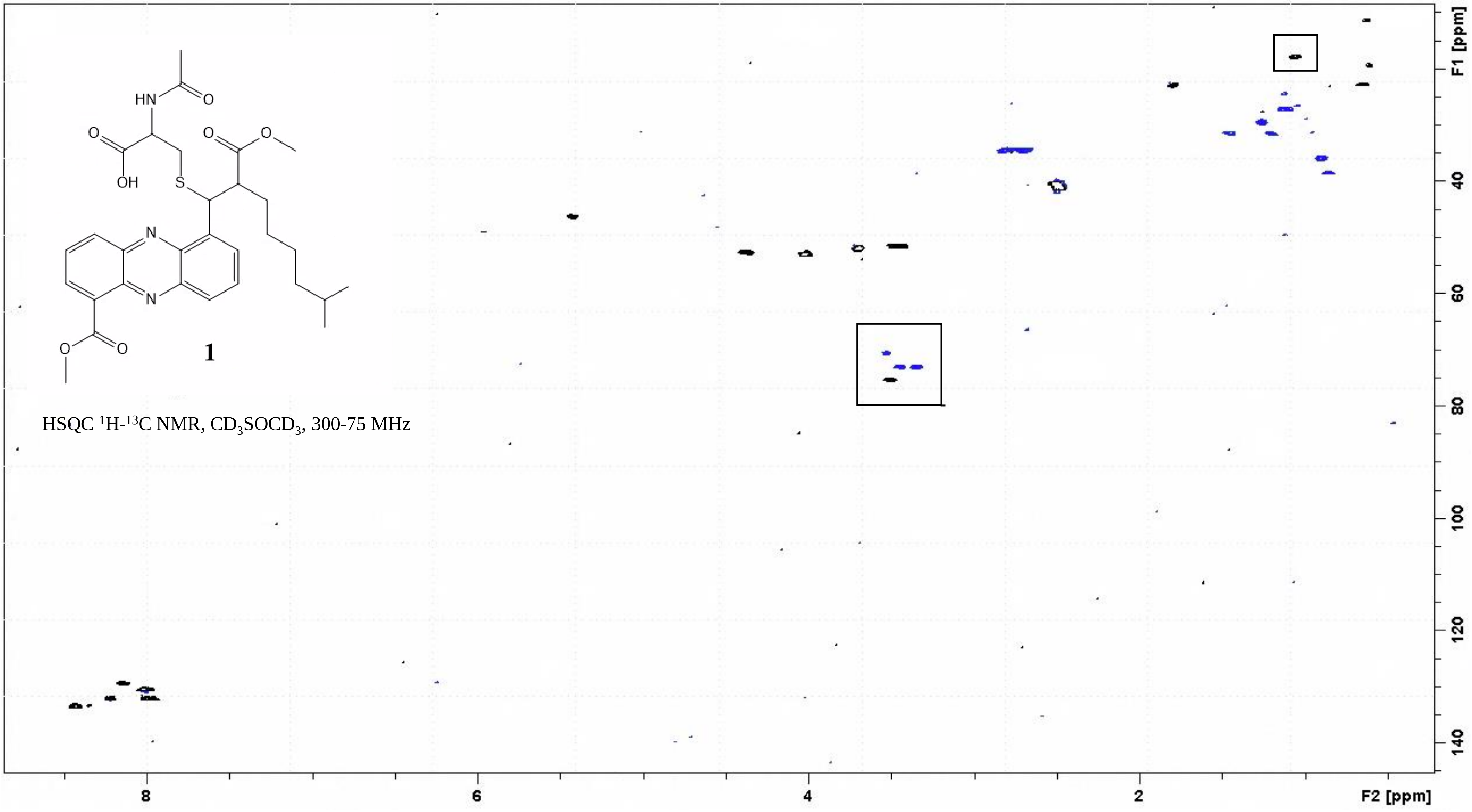


Supplementary Figure XII: HSQC of **1** in dmso-$d_6$ at 350K / 77°C, in the boxes PPG impurities signals

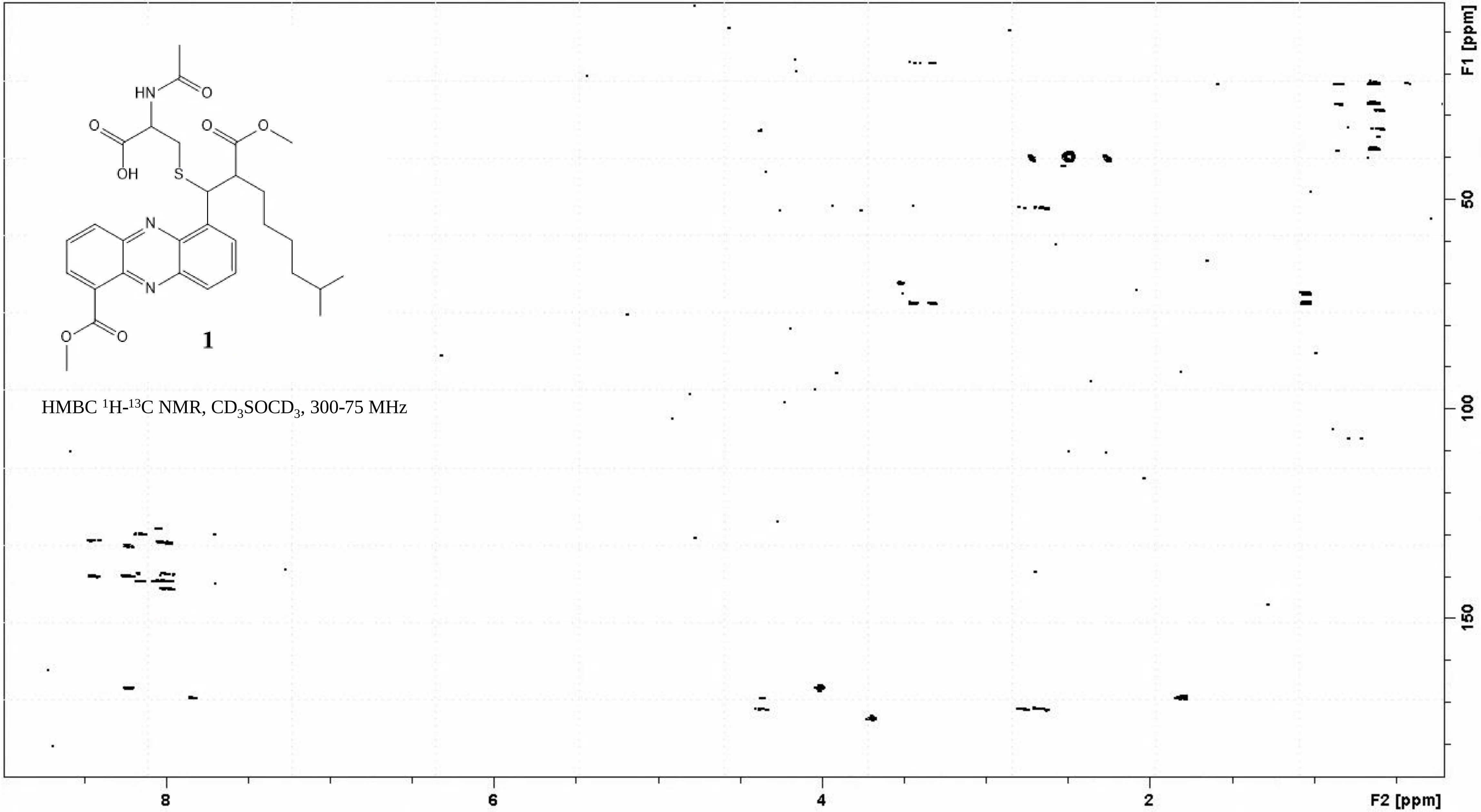


Supplementary Figure XIII: HMBC of **1** in dmso-$d_6$ at 350K / 77°C

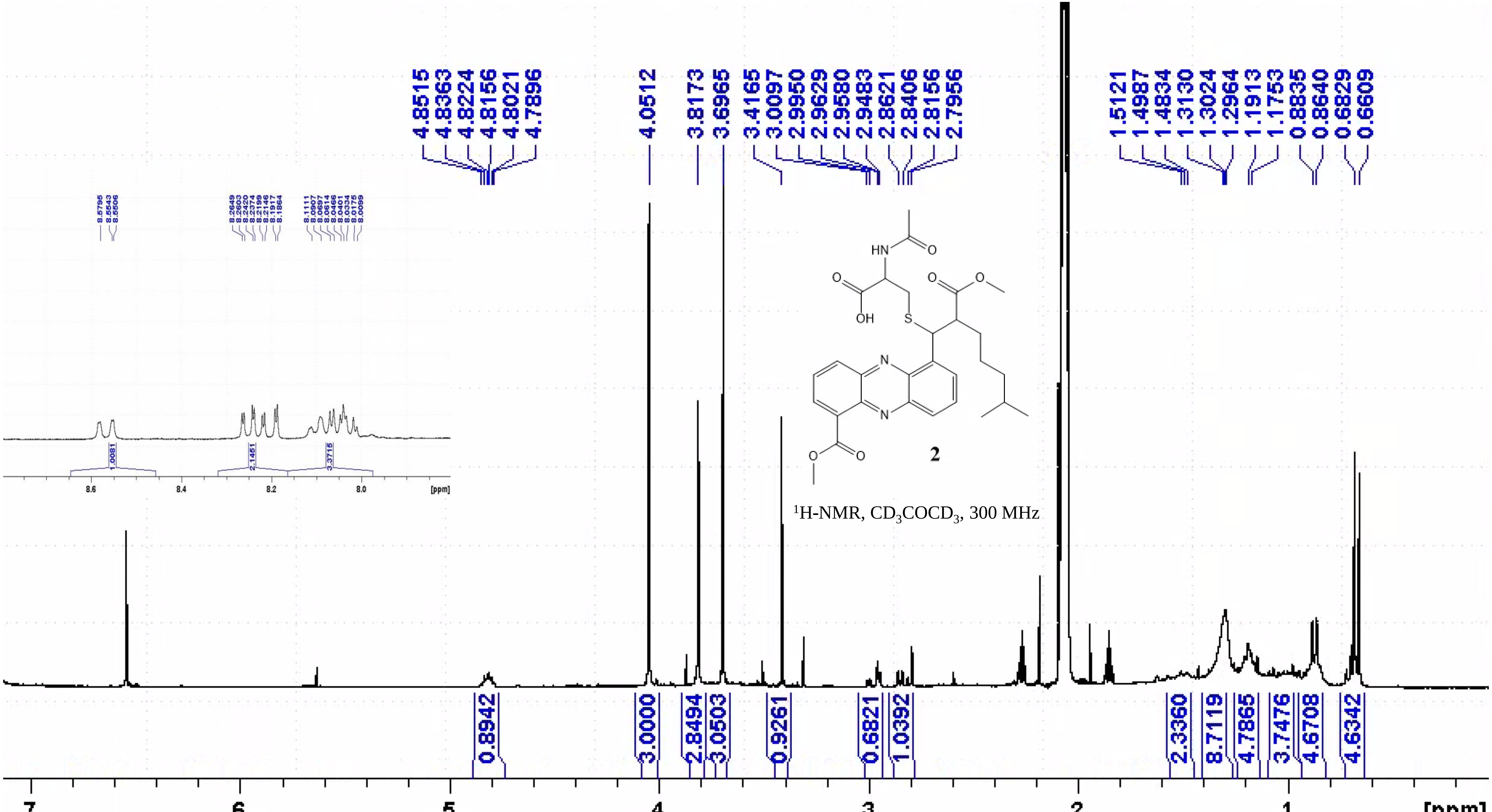


Supplementary Figure XIV: $^{1}$H-NMR of **2** in acetone-$d_6$ at 300K

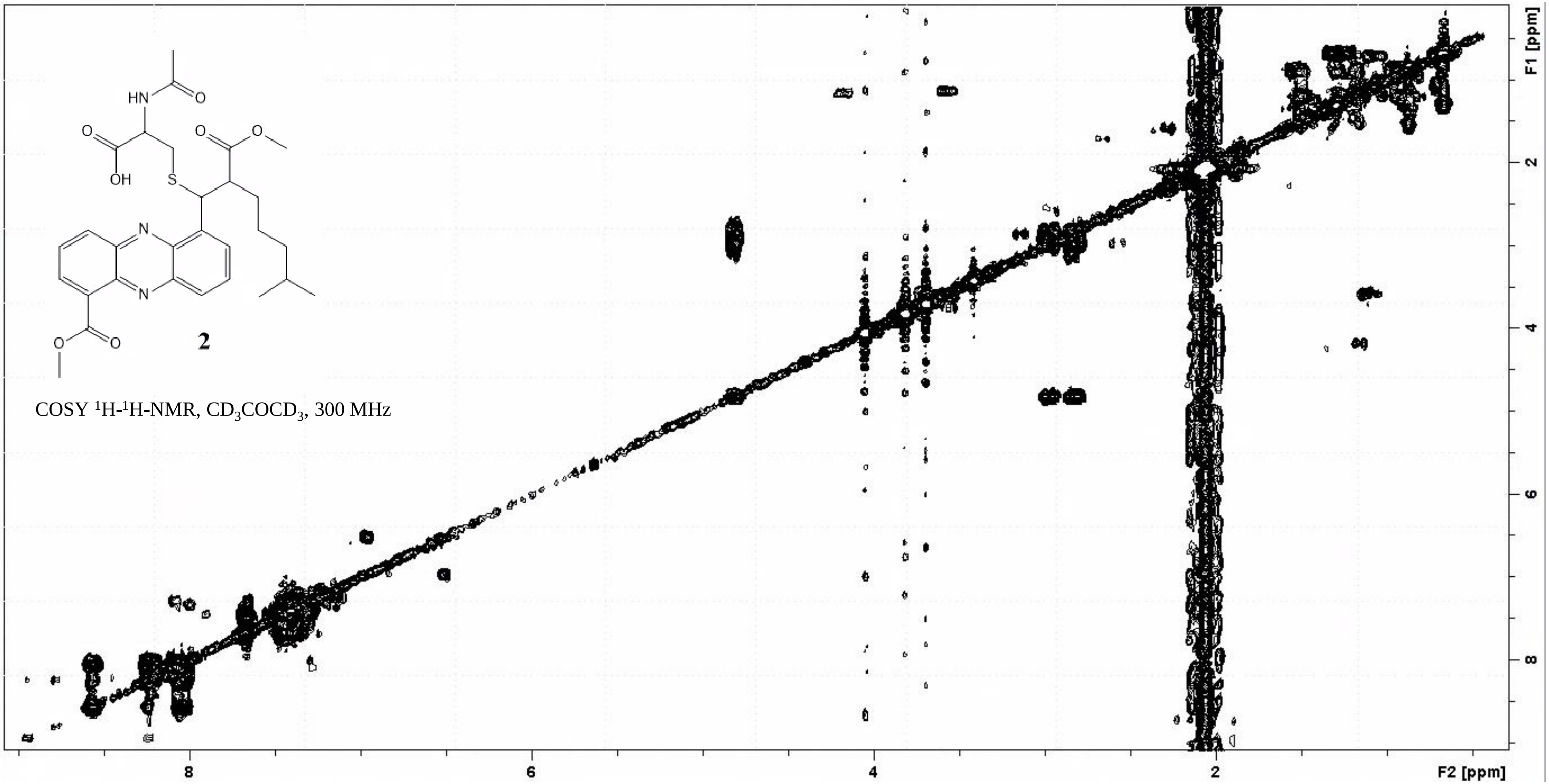


Supplementary Figure XV: COSY of **2** in acetone-$d_6$ at 300K

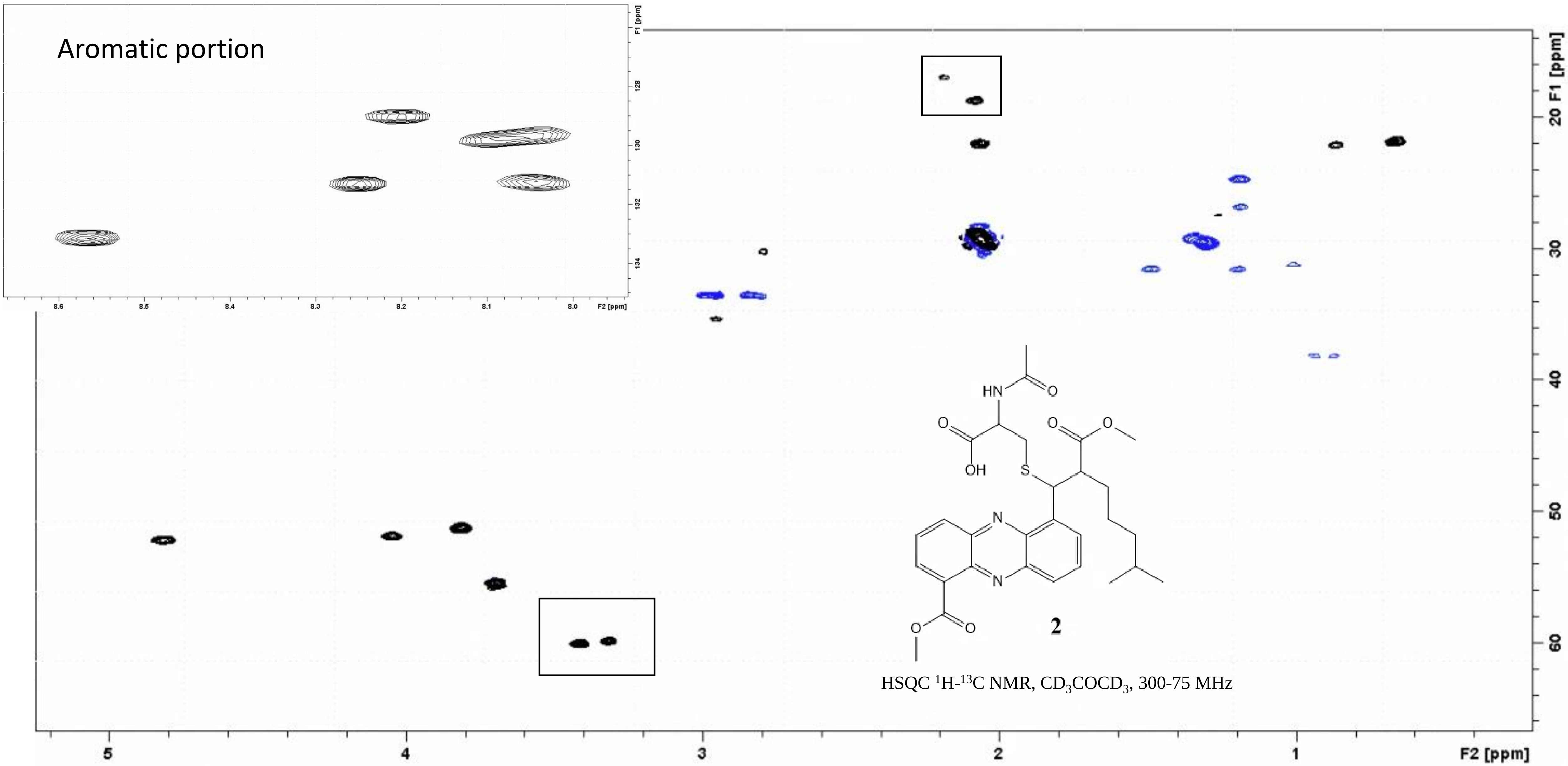


Supplementary Figure XVI: HSQC of **2** in acetone-$d_6$ at 300K, in the boxes signals from unrelated impurities

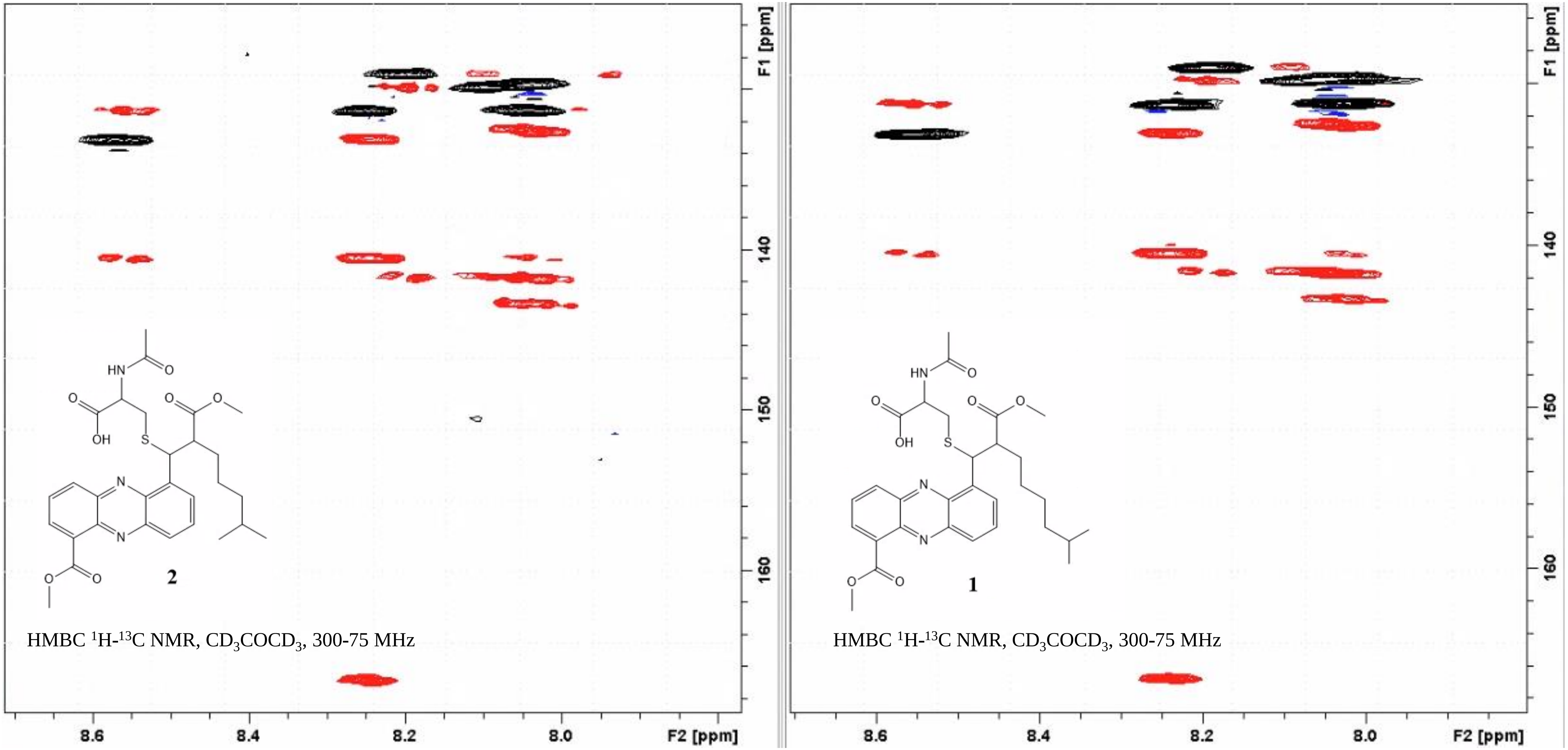


Supplementary Figure XVII: HSQC (black) and HMBC (red) for **2** (left) and **1** (right) in acetone-$d_6$ at 300K: aromatic portion

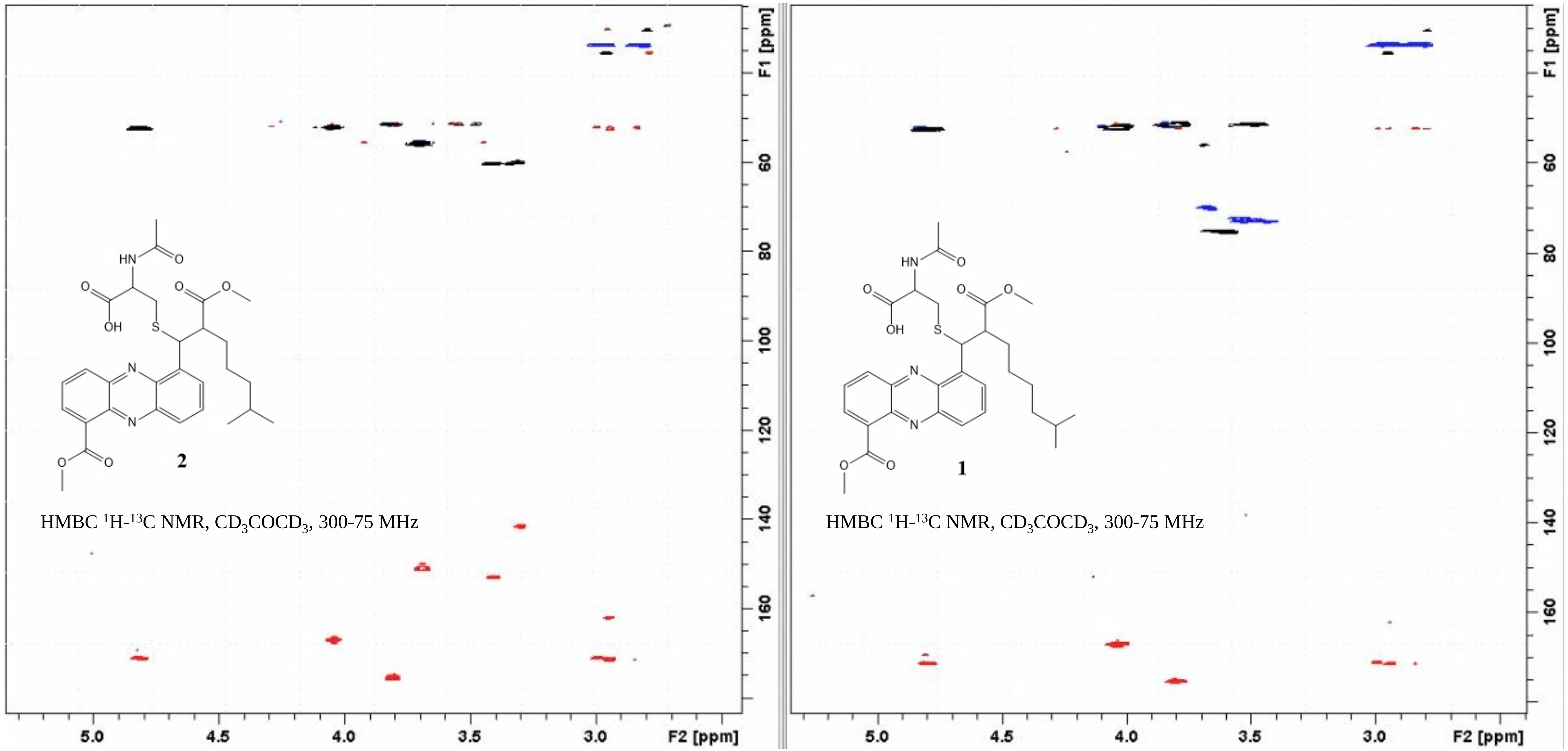


Supplementary Figure XVIII: HSQC (black) and HMBC (red) for **2** (left) and **1** (right) in acetone-$d_6$ at 300K: central portion

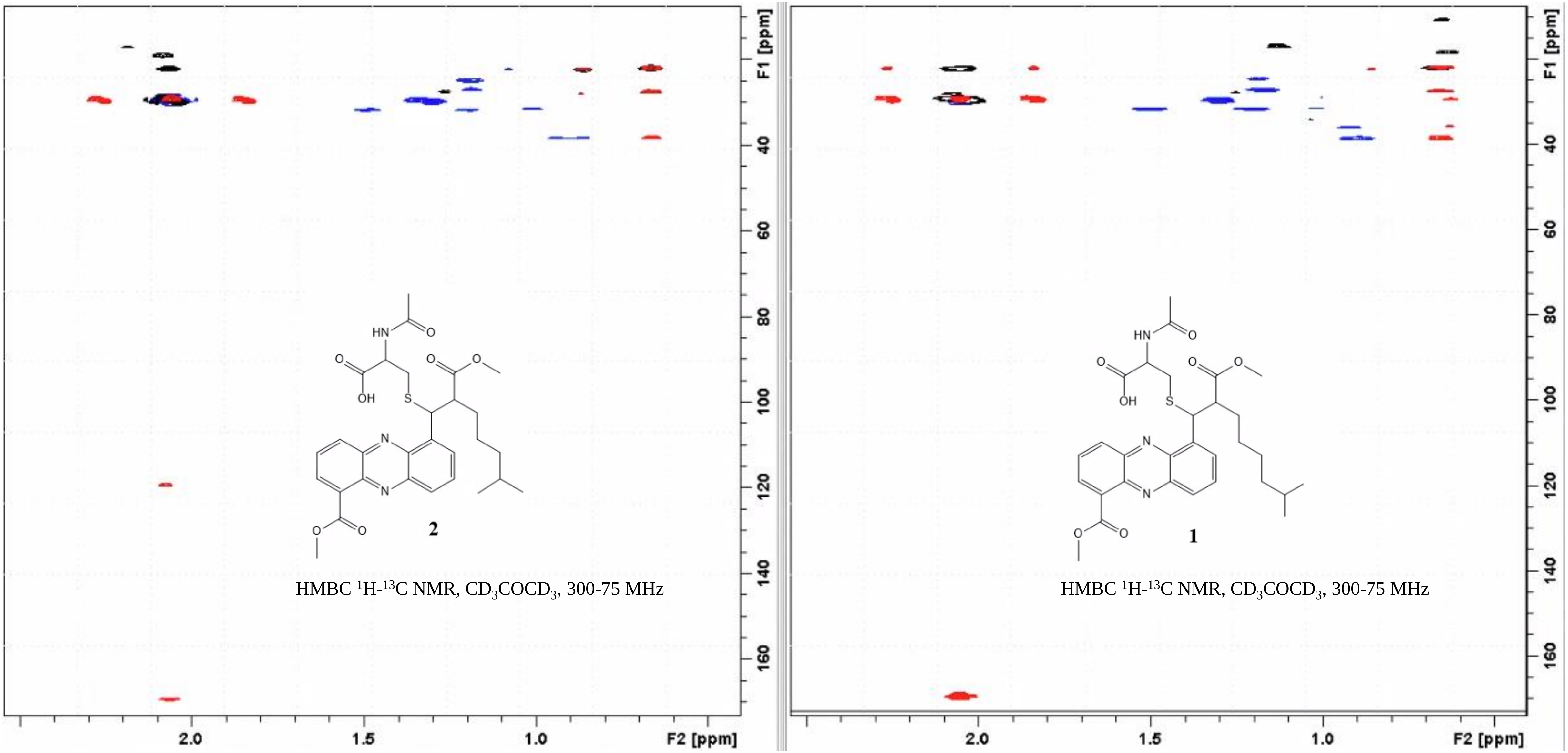


Supplementary Figure XIX: HSQC (black) and HMBC (red) for **2** (left) and **1** (right) in acetone-$d_6$ at 300K: aliphatic portion
